# Airborne Nanoplastics Perturb Mitochondrial Complex I via the ND6 Axis: Polymer-Specific Mitoepigenetic Remodeling Integrating Experimental, In Silico, and Machine Learning Analyses

**DOI:** 10.64898/2025.12.19.695405

**Authors:** Pradyumna Kumar Mishra, Aneha K Rajan, Apoorva Chouksey, Vikas Gurjar, Aniket Aglawe, Ashwani Pathak, Rajnarayan Tiwari, Devojit Kumar Sarma, Ravi Prakash Tiwari, Rupesh K. Srivasatava

## Abstract

Airborne nanoplastics constitute an emerging class of environmental contaminants, but their mitoepigenetic effects on human immune cells have not been systematically investigated. Ex vivo human lymphocytes were used to investigate integrated mitochondrial, epigenetic, and inflammatory responses induced by polystyrene (PS), polypropylene (PP), and polyvinyl chloride (PVC) nanoplastics. Fluorescence microscopy at multiple exposure time points and flow cytometry confirmed efficient cellular internalization and progressive intracellular accumulation of nanoplastics. Exposure elicited coordinated transcriptional remodeling of genes regulating mitochondrial dynamics (DRP1, MFN1), mitochondrial DNA encoded oxidative phosphorylation components (MT-ATP6, MT-COX1, MT-ND6), DNA repair (OGG1, APE1), DNA methylation machinery (DNMT1, DNMT3a, DNMT3b), and mitochondrial-associated miRNAs (miR-21, miR-34a, miR-155). Functional analyses revealed polymer and time-dependent disruption of mitochondrial membrane potential and respiratory chain activities, with Complex I identified as the primary site of vulnerability. Correlation analysis showed strong positive associations among DRP1, OMA1, DELE1, and ND6 (r > 0.9, R² > 0.8, p < 0.001), reflecting coordinated mitochondrial stress and epigenetic signaling, while negative correlations between DRP1 and MFN1 (r = -0.54, R² = 0.29, p < 0.01) and between APE and ND6 (r ≈ -0.42, R² ≈ 0.18, p < 0.05) highlight antagonistic regulation and impaired mitochondrial network stability linked to Complex I dysfunction. In silico docking of oxidized nanoplastic oligomers identified high-affinity interactions at the Complex I Fe-S cluster and cofactor-binding sites, suggesting direct interference with electron transfer. A random forest-based model accurately predicted MT-ND6 expression from Complex I activity (R² > 0.85), establishing a data-driven Complex I-ND6 axis. Collectively, these findings demonstrate that airborne nanoplastics induce integrated mitoepigenetic and immunometabolic dysregulation, underpinned by coordinated and antagonistic regulatory interactions in lymphocytes.

## Introduction

Plastics, integral to global industries, degrade into microplastics (<5 mm) and nanoplastics (NPs) (<1000 nm), resulting in pervasive environmental contamination and human exposure with emerging evidence of significant toxicological risks (WHO, 2024). Due to their nanoscale size, stability, and reactivity, NPs such as polystyrene (PS), polypropylene (PP), and polyvinyl chloride (PVC) persist in the environment and pose growing ecological and health risks (Enfrin et al., 2019; Nazeer et al., 2024). Since their atmospheric transport was first reported in 2015, they have subsequently been recognized as globally long-range distributed airborne pollutants (Dris et al., 2015). Their primary exposure routes in humans are inhalation, allowing the NPs to enter and accumulate throughout the body (Prata, 2018; Wang et al., 2024; Ratre et al., 2024). Upon exposure, they cross biological barriers and acquire a dynamic ‘protein-lipid corona’ that dictates their biological identity, cellular uptake, and toxicological profile (Lima et al., 2020). Once NPs enter the cellular environment, they initially bind to the plasma membrane (Lesniak et al., 2013), followed by internalization through either passive transmembrane diffusion or active endocytic mechanisms (Fiorentino et al., 2014).

Emerging evidence indicates that mitochondrial epigenetic regulation shaped by redox state, metabolic intermediates, and non-coding RNA (ncRNA) signaling constitutes a sensitive yet underexplored target of environmental stressors (Lees et al., 2023; Rajan et al., 2025). NPs may alter mitochondrial gene regulation by disrupting redox balance and metabolic flux, thereby modulating mtDNA-encoded transcriptional programs independently of overt structural damage (Tao et al., 2024; Mondal et al., 2025). Such effects are plausibly associated with NPs-induced impairment of electron transport chain (ETC) activity, enhanced reactive oxygen species (ROS) generation, and dysregulated calcium homeostasis, which together may contribute to sustained bioenergetic and immunometabolic dysfunction (Bu et al., 2024; Chulkov et al., 2025). In murine models, such exposure has been linked to inflammatory responses, immunotoxic effects, and neurological dysfunction (Wang et al., 2021; Prosperi et al., 2025), whereas in vitro studies across diverse human cell lines confirm efficient NPs internalization (Sendra et al., 2019; Xu et al., 2021), suggesting that damage to sensitive subcellular organelles, particularly mitochondria, represents a central mechanism underlying NPs-induced cellular toxicity.

Airborne NPs represent a clinically relevant immunotoxic exposure route, as inhaled particles can enter systemic circulation and directly engage immune cells. Peripheral blood mononuclear cells (PBMCs) were selected as our ex vivo model due to their high mitochondrial content and sensitivity to environmental stressors, allowing effective reflection of systemic immune and metabolic perturbations (Soni et al., 2025a; Mishra et al., 2022). These characteristics make PBMCs particularly suitable for evaluating NPs-induced mitochondrial dysfunction and associated epigenetic alterations (Hernández et al., 2025; Garrafa et al., 2023). PS, PP, and PVC were chosen for comparative analysis based on their environmental prevalence, frequent detection in airborne pollutants (Visileanu et al., 2025), and documented presence in human tissues, including lung, blood, placenta, and brain (Thompson et al., 2024), that may promote oxidative stress and mitochondrial dysfunction, characterized by excessive ROS production, impairment of ETC complexes, mitochondrial membrane depolarization (ΔΨm), ATP depletion, and mtDNA damage. These mitochondrial disturbances are tightly coupled to inflammatory signaling and pro-inflammatory cytokines (Bhargava et al., 2019).

In this study, we investigated the effects of PS, PP, and PVC on mitochondrial dysfunction and epigenetic regulation in PBMCs, providing a comprehensive framework to elucidate NPs-induced immunometabolic dysfunction and associated long-term risks of chronic disease. A key focus is understanding how NPs alter mitochondrial gene expression, particularly the mtDNA-encoded ND6 gene, a critical subunit of complex I that regulates electron flux and complex stability within this major redox and ROS-generating hub of OXPHOS. Perturbations in ND6 can exacerbate electron leakage, amplify oxidative stress, and trigger retrograde stress signaling; despite its mechanistic importance, its role in NPs-induced mitochondrial dysfunction and epigenetic regulation has not been systematically explored until now. To address this gap, we employed novel molecular docking simulations to estimate direct NPs interactions with OXPHOS component Complex I, revealing potential binding sites and their impacts on electron transport, ATP synthesis, and mitochondrial homeostasis. Furthermore, we applied a machine learning-based supervised multi-output regression framework to model relationships between experimentally measured Complex I activity and PCR-quantified ND6 expression, uncovering AI-driven predictive patterns that link enzymatic function to gene regulation and complement traditional assays for enhanced mechanistic insights. By integrating these experimental, in silico, and AI-assisted approaches, this work establishes a novel multidisciplinary strategy to clarify how NPs exposure in PBMCs disrupts mitochondrial and epigenetic integrity.

## 2. Materials and methods

### 2.1 Blood sample collection and lymphocyte isolation

Peripheral blood samples were obtained by sterile venipuncture into heparinized tubes following approval from the Institutional Ethics Committee (IEC) of the ICMR-National Institute for Research in Environmental Health (ICMR-NIREH). Samples were transported on ice (2-8°C) and processed within 2-3 h of collection. PBMCs were isolated using HiSep™ LSM 1077 (HiMedia, Mumbai, India), washed, and resuspended in 1X PBS (Cell Signaling Technology Danvers, MA, USA). The isolated PBMCs were then stored at -20°C until further analysis (Bhargava et al., 2015; Bunkar et al., 2020).

### 2.2 Characterization and uptake of NPs

The particle size distribution, aggregation behavior, and surface charge (zeta potential) of the NPs were assessed using the Malvern Panalytical Zetasizer Nano ZS (Malvern, Worcestershire, UK) equipped with disposable cuvette-based measuring cells. For all measurements, NPs suspensions were prepared in distilled water and gently vortexed to ensure homogeneous dispersion. For size determination, dynamic light scattering (DLS) was performed, providing measurements of the hydrodynamic diameter and polydispersity index (PDI). Surface charge analysis was carried out via electrophoretic light scattering (ELS) to determine the zeta potential (Gurjar et al., 2024; Arranz et al., 2024). Each sample was equilibrated at 25°C prior to measurement, and all readings were acquired in triplicate to ensure accuracy and reproducibility. In vitro cellular uptake of nanoplastics was evaluated by following a previously described protocol using fluorescently labeled polystyrene nanoplastics (PS-NPs) (FluoSpheres™ carboxylate-modified microspheres, Thermo Fischer Scientific, Waltham, MA, USA). Lymphocytes were exposed to PS-NPs under standard culture conditions and collected at 0, 30, 60, and 120 min to assess time-dependent uptake. Qualitative visualization of NPs uptake was performed using fluorescence microscopy (IX83, Olympus Corporation, Shinjuku-ku, Tokyo, Japan) after thorough washing with phosphate-buffered saline (PBS) to remove unbound particles, while quantitative analysis of uptake kinetics was carried out by flow cytometry using an Attune™ NxT flow cytometer (Thermo Fischer Scientific, Waltham, MA, USA), measuring fluorescence intensity within gated lymphocyte populations (Bhargava et al., 2018).

### 2.3 Treatment and incubation of lymphocytes with NPs

Stock suspensions of PS, PP, and PVC were prepared at a concentration of 1 µM each. For exposure experiments, 2.5 mL of RPMI-1640 medium was dispensed into sterile Petri dishes and supplemented with 100 µL of washed PBMCs. The cultures were maintained in a CO₂ incubator, and samples were collected at 6, 24, and 48 h post-exposure. Cells were harvested by centrifugation at 4,500 × g for 10 min, resuspended in PBS, and stored at -20°C until analysis. Cell-free supernatants were collected separately and stored under the same conditions

### 2.4 Analysis of cellular oxidative damage and intracellular stress

Whole-cell genomic DNA was extracted using the PureLink Genomic DNA Mini Kit (Invitrogen, Thermo Fischer Scientific, Waltham, MA, USA) following the manufacturer’s protocol. Oxidative DNA damage was quantified by detecting oxidized purine bases, primarily deoxyguanosine, via formamidopyrimidine glycosylase (FPG) digestion with appropriate negative controls. Intracellular and mitochondrial reactive oxygen species (mtROS), apoptosis, and mitochondrial membrane potential (ΔΨm) were assessed to evaluate NPs-induced oxidative and cytotoxic effects. Total intracellular ROS were measured using CM-H₂DCFDA labeling in cells exposed to PS, PP, and PVC NPs at 1X, 10X, and 100X concentrations for 6, 24, and 48 h; fluorescence intensity was quantified fluorometrically. Mitochondrial ROS (mtROS) were assessed using the CellROX™ Deep Red Reagent (Thermo Fisher Scientific, Waltham, MA, USA), with cells stained at 500 nM for 1 h at 37°C and analyzed via flow cytometry and fluorescence microscopy. Apoptosis and necrosis were simultaneously evaluated using the CellEvent™ Caspase-3/7 Green Detection Reagent (Thermo Fisher Scientific, Waltham, MA, USA) with SYTOX™ AADvanced™ staining and flow cytometric analysis (488 nm excitation; 530/30 BP and 690/50 BP filters). ΔΨm was monitored to determine concentration and time-dependent mitochondrial depolarization (Bunkar et al., 2020).

### 2.5 Gene expression analysis by quantitative real-time PCR

Total RNA was isolated using TRIzol Reagent (Life Technologies, Thermo Fischer Scientific, Waltham, MA, USA) following the manufacturer’s protocol. Gene expression profiling was performed to evaluate mitochondrial dynamics (fission: DRP1; fusion: MFN1) and mitochondrial genome-encoded transcripts (MT-ATP6, MT-COX1, MT-ND6). Additionally, genes involved in the integrated stress response (OMA1, DELE1), DNA damage repair (APE1, OGG1), and DNA methylation (DNMT1, DNMT3a, DNMT3b) were quantified using quantitative real-time PCR (qRT-PCR). Cycle threshold (Ct) values were obtained, and relative expression levels were calculated using the 2^^−ΔΔCt^ method. GAPDH served as the internal normalization control (Soni et al., 2025a).

### 2.6 Quantification of DNA methylation

mtDNA methylation analysis was performed using bisulfite conversion followed by methylation-specific PCR (MSP). Genomic DNA was isolated from NPs-exposed cells (PS, PP, and PVC; 6, 24, and 48 h) using the QIAamp DNA Blood Mini Kit (QIAGEN, Hilden, Germany). A total of 3000 ng DNA was digested with BamHI at 37°C for 4 h. Following digestion, 15 µL of DNA was mixed with 125 µL bisulfite reagent, incubated at 98°C for 10 min and 60°C for 150 min, and subsequently desulfonated, purified, and eluted. MSP was performed using four primer sets targeting the D-loop1, D-loop2, 12S rRNA, and 16S rRNA regions. PCR reactions (20 µL) contained 50 ng bisulfite-converted DNA using the Luna® Universal qPCR Kit (New England Biolabs, Ipswich, MA, USA). Amplification was carried out on the Insta Q-96™ real-time PCR system (HiMedia Laboratories, Mumbai, India). Ct values were obtained at the end of cycling and reported as mean Ct for each mitochondrial target region (Mishra et al., 2022a).

### 2.7 Gene expression analysis of miRNA and its target genes

Quantitative RT-PCR was performed to assess the expression of miR-21, miR-34a, and miR-155, key regulators of inflammation, apoptosis, and oxidative stress. Plasma miRNA was isolated using the PureLink™ miRNA Isolation Kit (Thermo Fischer Scientific, Waltham, MA, USA) and reverse-transcribed into cDNA using miRNA-specific primers. PCR amplification was carried out under optimized thermal cycling conditions, and products were resolved on HiMedia Agarose (HiMedia Laboratories, Mumbai, India) stained with SYBR® Safe (Invitrogen, Thermo Fischer Scientific, Waltham, MA, USA) for UV visualization. Expression of miRNA target genes PTEN, FOXO3, and Bcl2 was quantified by qRT-PCR using GAPDH as an internal control. Relative expression levels were calculated using the 2^-ΔΔCt method (Mishra et al., 2021).

### 2.8 Evaluation of inflammatory cytokine levels

To evaluate the inflammatory response, the levels of Interleukin-6 (IL-6), Interleukin-8 (IL-8), and Tumor Necrosis Factor (TNF-α) were quantified using Human GENLISA™ ELISA kits (KRISHGEN BioSystems, Cerritos, CA, USA), following the manufacturer’s instructions. Absorbance was measured using the Spark® multimode microplate reader (Tecan, Switzerland) (Mishra et al., 2022b).

### 2.9 Assessment of mitochondrial electron transport chain complex activity

Cells were harvested after NPs treatment at 6, 24, and 48 h, and assays were performed according to the manufacturer’s protocols. Further processed for studying mitochondrial ETC activity using MitoCheck Complex Activity Assay Kits (Cayman Chemical, Ann Arbor, MI, USA), and absorbance was measured using the Spark® multimode microplate reader (Tecan, Switzerland) (Soni et al., 2025a).

### 2.10 Statistical analysis

All statistical analyses were performed using Python. Data processing and organization were conducted with Pandas and NumPy, while Matplotlib and Seaborn were used for graphical visualization of gene expression and biochemical variations. Data normalization was carried out using the StandardScaler function from Scikit-learn (sklearn), followed by principal component analysis (PCA) to identify major sources of variance and K-means clustering to classify treatment groups based on expression profiles. Statistical comparisons between control and NPs-exposed groups were evaluated using the SciPy stats module (Kaur et al., 2025).

### 2.11 In silico docking studies

Molecular docking was performed to investigate the interactions between oxidatively functionalized NPs oligomers and key mitochondrial proteins. The crystal structure of mitochondrial complex I (PDB ID: 5XTD) was retrieved from the Protein Data Bank and prepared using Maestro’s Protein Preparation Wizard (Schrödinger Suite). Protein preparation included adding hydrogens, assigning bond orders, optimizing protonation states, refining the hydrogen-bond network, and restrained minimization under the OPLS4 force field. Water molecules distant from active sites were removed to improve ligand accessibility, and receptor grids were generated around functionally relevant binding regions. A library of sixteen oxidized NPs oligomers derived from PS, PP, and PVC, functionalized with -OH, -COOH, or -CHO groups at α- or ω-termini, was constructed and optimized using LigPrep at physiological pH (7.4 ± 0.2). Docking simulations were performed with Glide in Extra Precision (XP) mode, generating up to ten poses per ligand, followed by post-docking minimization. Binding poses were ranked by GlideScore (kcal/mol), with greater scores indicating stronger interactions within catalytic or cofactor-binding domains. Protein-ligand interactions were visualized and analyzed in Maestro’s Pose Viewer to identify hydrogen bonds, hydrophobic contacts, electrostatic interactions, and π-π stacking. All computational steps followed validated Schrödinger workflows to ensure reproducibility and structural accuracy (Soni et al., 2025b).

### 2.12 AI prediction of ND6 expression from mitocomplex I activity

The AI-based prediction of ND6 gene expression from Mitocomplex 1 activity was performed by constructing a multi-output regression model. The activity of Mitocomplex 1 was experimentally measured using an ELISA assay, while ND6 gene concentration was accurately quantified following PCR amplification. For the machine learning analysis, the measured variables representing Mitocomplex 1 activity were used as input features, and the PCR-quantified ND6 gene expression values were used as output targets. All data variables underwent normalization using standard scaling to facilitate efficient learning by the model. The dataset was randomly split into training (80%) and test (20%) subsets to ensure objective assessment of predictive performance. The model implementation used a Random Forest Regressor wrapped in a Multioutput Regressor, enabling simultaneous prediction of multiple ND6 gene expression endpoints. After training on normalized data, generalization performance was quantified using R-squared and mean squared error metrics on the held-out test set. Residual and actual-versus-predicted plots were generated to visually assess model fit and error distributions, supporting robust evaluation of prediction quality (Kaur et al., 2025, Gurjar et al., 2024).

## 3. Results

### 3.1 Characterization and cellular uptake of NPs

Dynamic light scattering (DLS) and zeta potential analyses together characterized the physicochemical properties of the polymer NPs in aqueous suspension. PS exhibited a Z-average hydrodynamic diameter of 87.4 nm **(Figure 1A)**, PP showed a larger Z-average size of 148.4 nm **(Figure 1B)**, and PVC displayed the largest Z-average diameter of 227.4 nm **(Figure 1C)**, with a dominant intensity peak. Zeta potential measurements revealed near-neutral surface charge for all materials, with values of −3.39 mV for PS, −0.457 mV for PP, and −0.317 mV for PVC, indicating limited electrostatic stabilization in aqueous media. Time-dependent uptake of PS particles by lymphocytic cells was first evaluated using DIC fluorescence microscopy. At baseline (0 min) **(Figure 2A)**, cells exhibited a uniform cytoplasmic appearance with no detectable fluorescent signal, indicating the absence of particle internalization. After 1 h of incubation **(Figure 2B)**, discrete punctate fluorescent signals were observed within the cells, consistent with the initial internalization of PS particles. With prolonged exposure, intracellular fluorescence increased markedly. At 3 h **(Figure 2C)**, numerous fluorescent puncta were distributed throughout the cytoplasm, suggesting active and sustained uptake. By 6 h **(Figure 2D)**, extensive intracellular accumulation of PS was evident, characterized by dense and widespread fluorescent signals occupying a substantial portion of the cytoplasmic area. To quantitatively validate the microscopy findings, flow cytometric analysis was performed using fluorescence intensity (BL1-H) as a measure of PS internalization. Overlaying staggered histograms revealed a progressive rightward shift in fluorescence intensity with increasing exposure time compared to the untreated control. The control population displayed a narrow fluorescence distribution within the low-intensity range, representing baseline autofluorescence. Cells analyzed at 0 min showed a fluorescence profile largely overlapping with the control, indicating minimal PS association. After 30 min of exposure, a discernible rightward shift in the fluorescence peak was observed, indicating the onset of particle uptake. This shift became more pronounced at 60 min, with a substantial increase in mean fluorescence intensity. Cells exposed for 120 min exhibited the greatest rightward displacement and highest fluorescence intensity, reflecting maximal PS uptake among the tested time points. **(Figure 3)**

**Figure 1:**
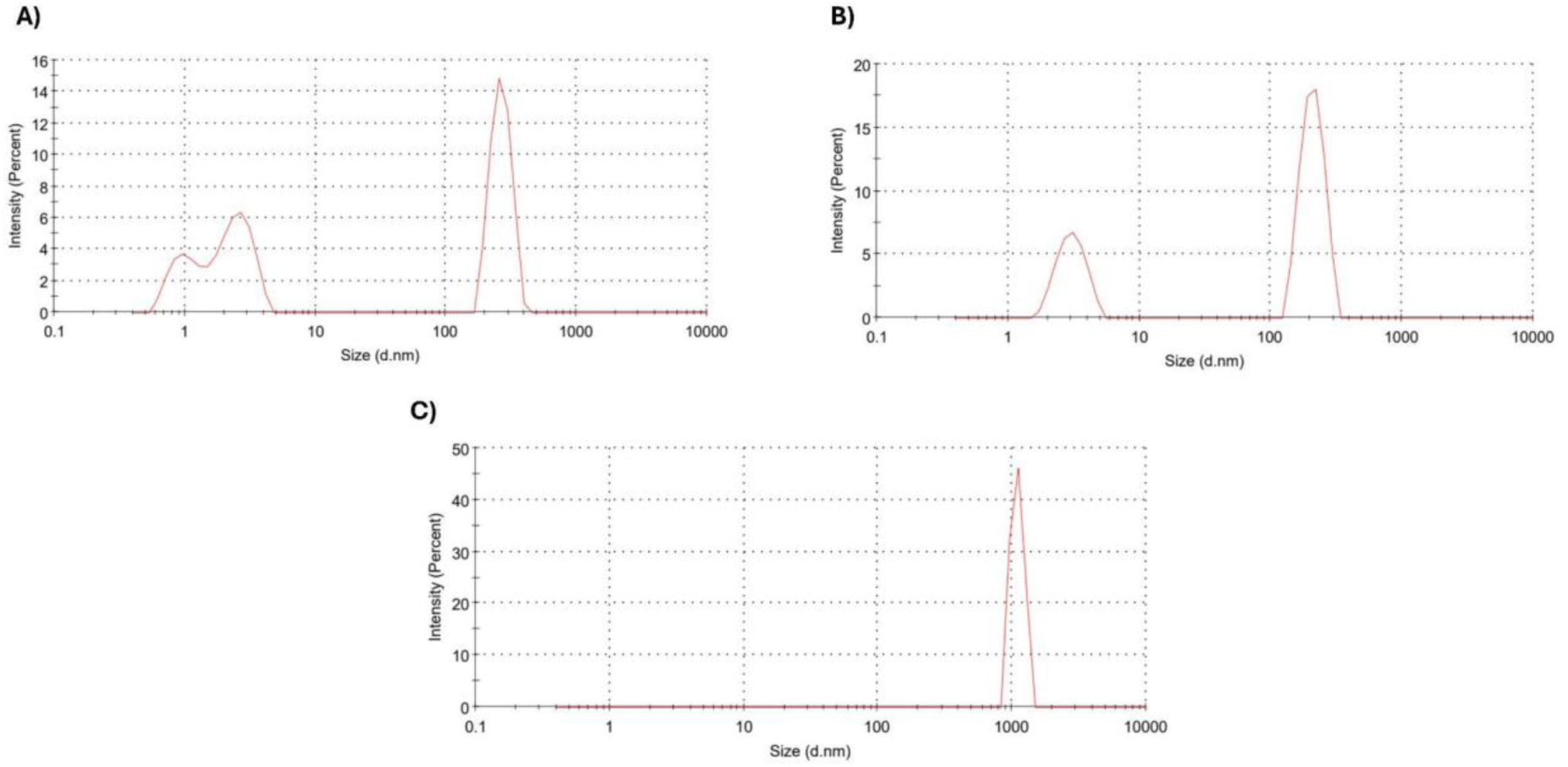
Particle size distribution of polymer NPs measured by dynamic light scattering (DLS). DLS analysis showing the hydrodynamic size distributions of **(A)** polystyrene (PS), **(B)** polypropylene (PP), and **(C)** polyvinyl chloride (PVC) NPs in aqueous suspension.

**Figure 2:**
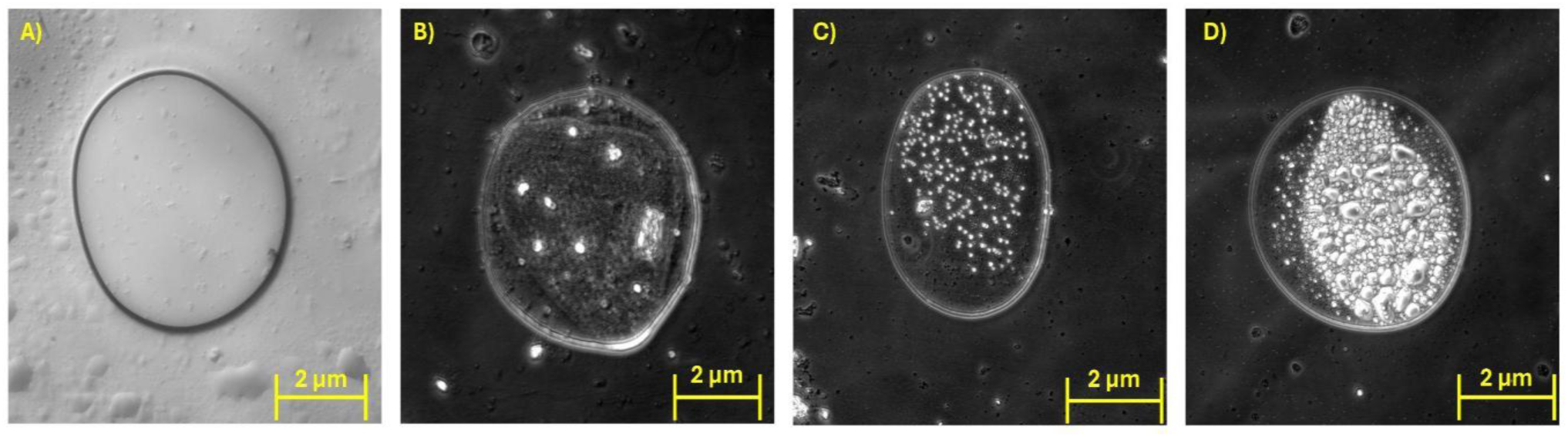
Time-dependent uptake of PS particles by lymphocytic cells visualized using DIC fluorescence microscopy. **(A)** 0 min, **(B)** 1 h, **(C)** 3 h, and **(D)** 6 h post-incubation. Progressive intracellular accumulation of PS particles is evident with increasing incubation time. Scale bar: 2 μm.

**Figure 3:**
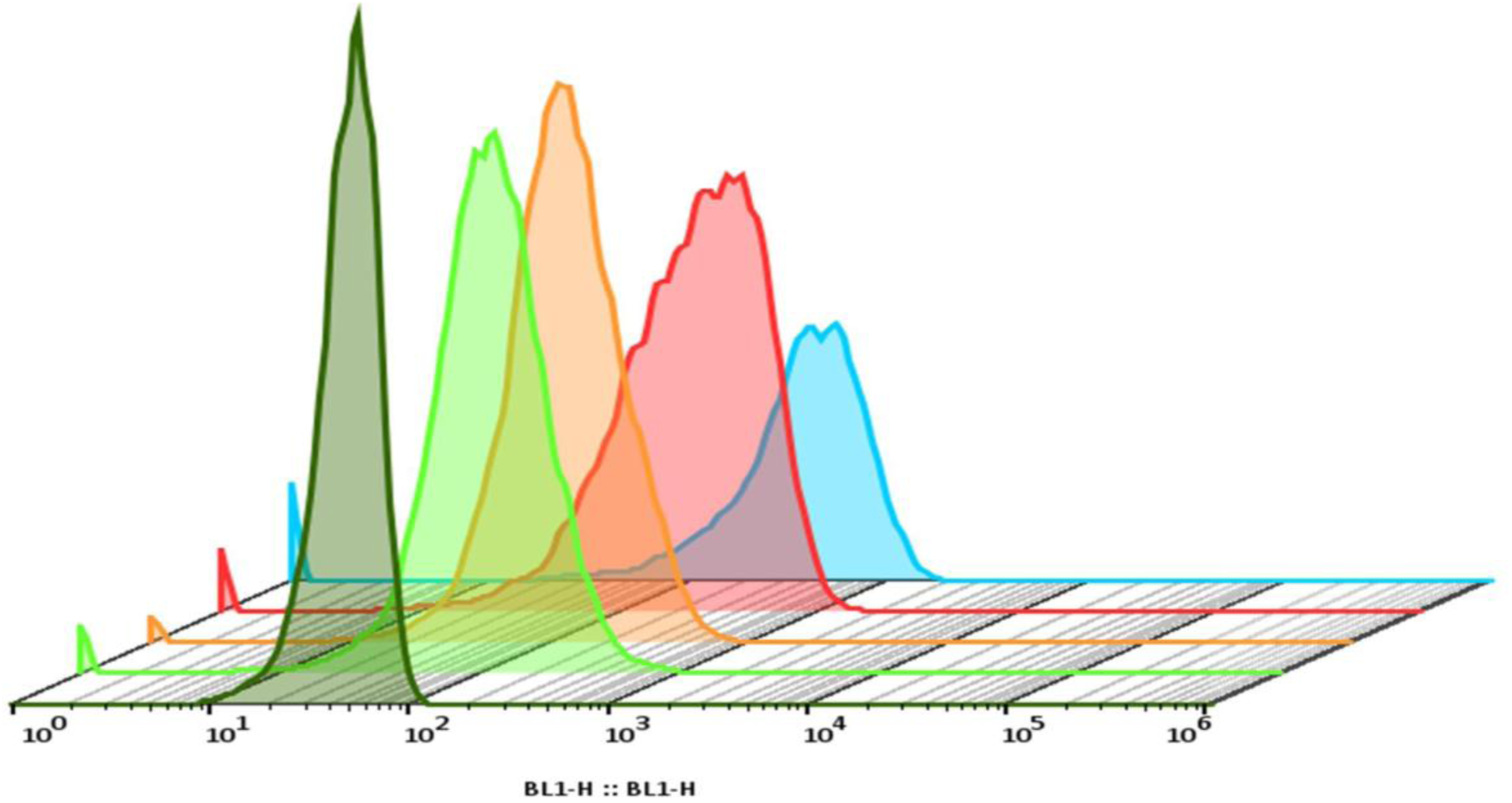
Flow cytometric analysis of time-dependent PS uptake by lymphocytic cells. Overlay staggered histograms show fluorescence intensity (BL1-H) of lymphocytic cells following exposure to PS particles at different time points. The untreated control is shown in black, 0 min in green, 30 min in orange, 60 min in red, and 120 min in blue. A progressive rightward shift in fluorescence intensity is observed with increasing exposure time, indicating enhanced cellular uptake of PS particles.

### 3.2 Oxidative damage and intracellular stress

The Fpg enzyme assay revealed a marked increase in oxidized purine bases in DNA from all NPs-treated groups relative to the control (**Figure 4)**. At 6 h, ROS levels were highest in PP, followed by PS and PVC. By 24 h, FAPY expression declined substantially across all exposure groups, although levels remained detectable, with PP and PVC exhibiting comparable intermediate activity. At 48 h, FAPY expression was further reduced, approaching minimal levels, particularly in PVC-exposed cells. In contrast, control samples displayed only baseline FAPY expression at 0 h, with no comparable induction observed. Flow cytometry using CM-H₂DCFDA (**Figure 5**) showed a rapid, concentration-dependent increase in intracellular ROS after PS, PP, and PVC exposure. PP elicited the strongest and most sustained ROS response, while PS and PVC produced transient early spikes followed by partial recovery and late-phase reactivation, indicating polymer-specific redox dynamics. mtROS assessed using CellROX™ Deep Red (**Figure 6**) confirmed these trends, showing maximal oxidative stress at 6 h and persistent elevation under PP exposure up to 48 h. Caspase-3/7 activation assays (**Figure 7**) revealed distinct, dose-dependent shifts in cell fate. Moderate doses (10X) induced the highest early and late apoptosis, particularly in PS- and PVC-treated cells at 24 h. At 100X, apoptotic signals declined while necrosis markedly increased, indicating a transition from programmed apoptosis to direct cytotoxic injury. PP triggered the most sustained necrotic response, aligning with its prolonged ROS output. ΔΨm analysis showed pronounced depolarization, with only ∼30-37% of cells retaining high ΔΨm at 6 h and further declines by 24 h, most severe in PVC-treated cells (∼13-17%) (**Figure 8**). These findings reflect cumulative mitochondrial dysfunction and progressive redox collapse.

**Figure 4:**
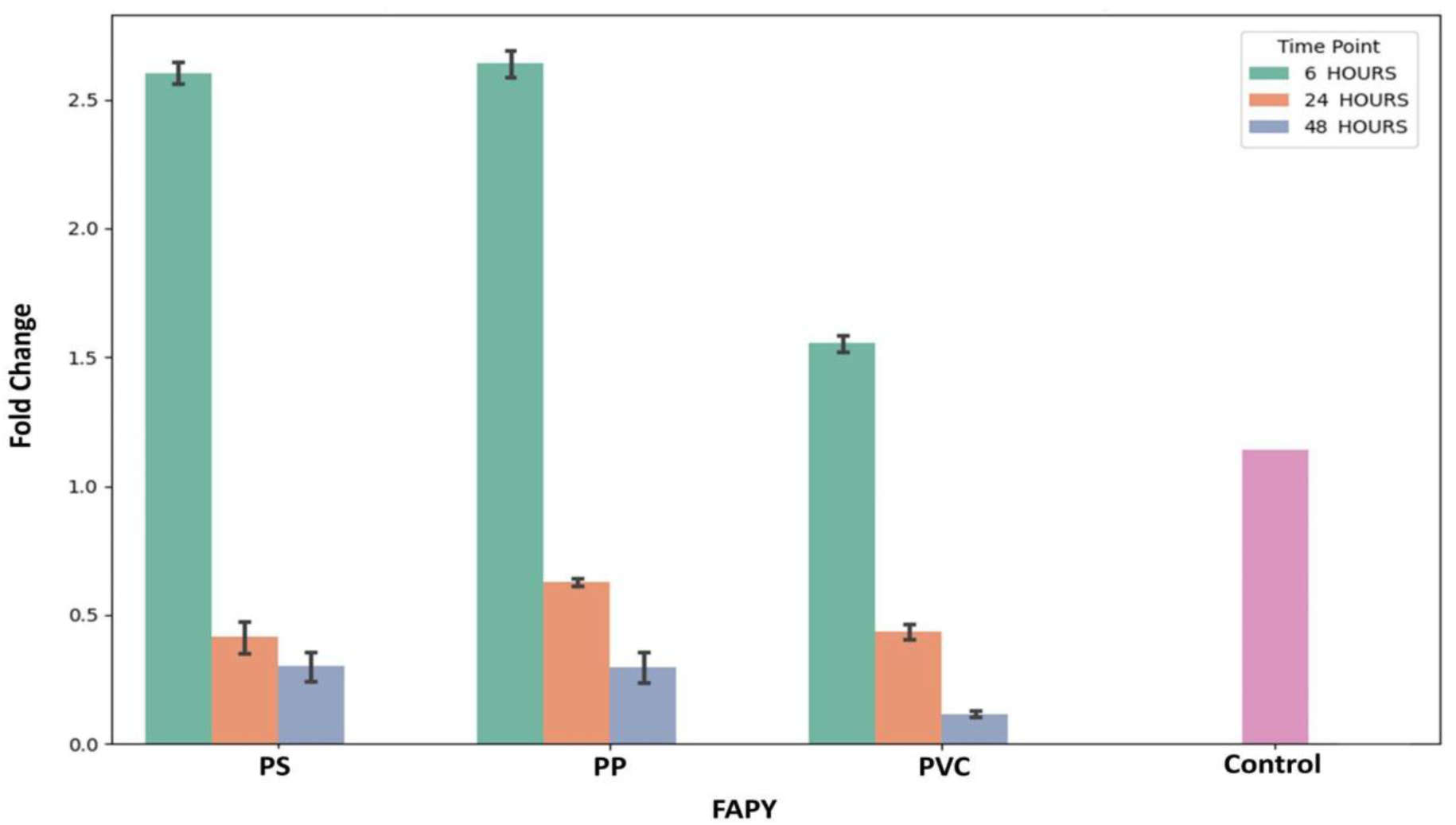
Bar graph showing the mean fold change (FC) in Fpg gene expression following exposure to PS, PP, and PVC for 6, 24, and 48 h compared to the untreated control (0 hr). A pronounced upregulation was observed at 6 h in all treatment groups, followed by a time-dependent decline at 24 and 48 h, indicating transient activation of the base excision repair (BER) pathway in response to NPs-induced oxidative stress. Error bars represent standard error of the mean (SEM).

**Figure 5:**
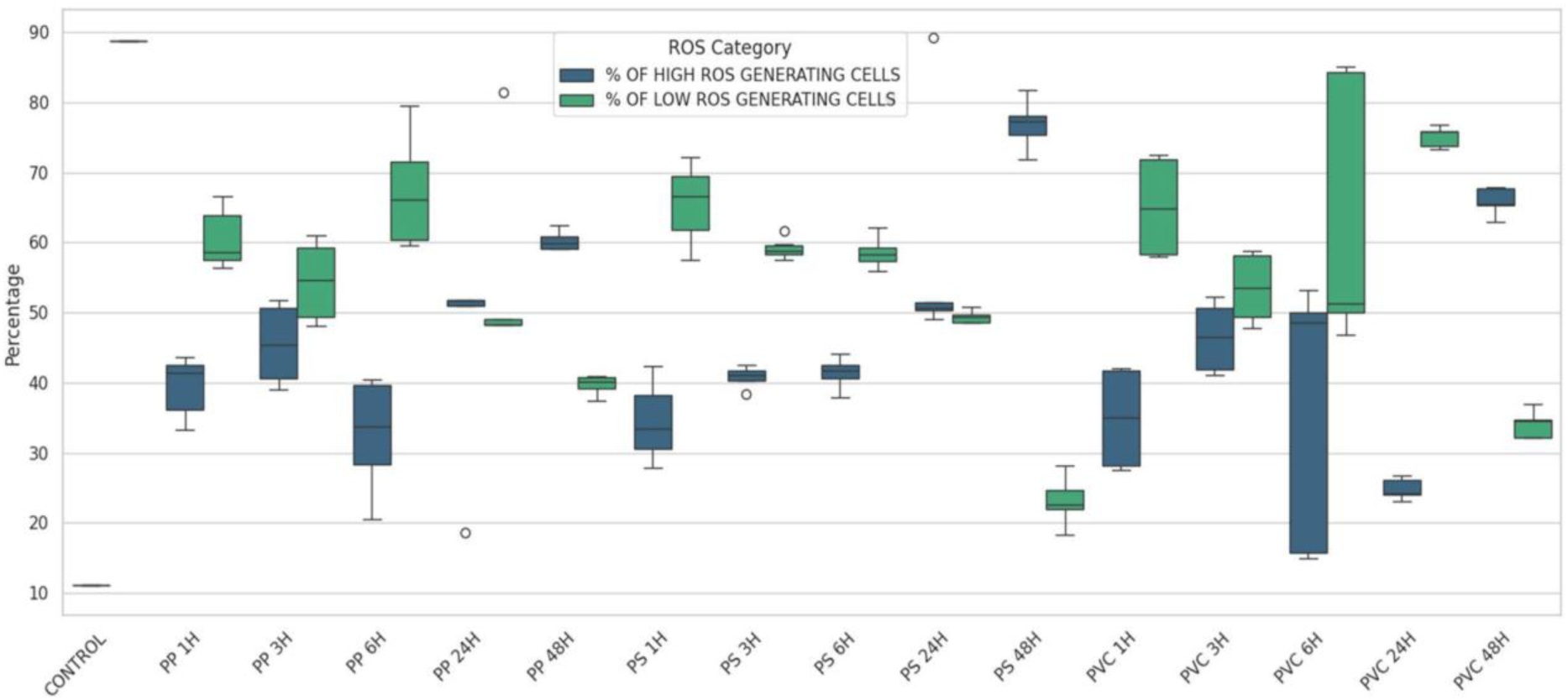
Distribution of the percentage of ROS-generating cells across different materials, exposure durations, and control conditions. This box-and-whisker plot depicts the variation in ROS generation among cells exposed to PS, PP, and PVC at different time points (1 h, 3 h, 6 h, 24 h, and 48 h), along with an untreated control. The y-axis represents the percentage of ROS-generating cells, while the x-axis (Material-Time) indicates each experimental condition. Dark blue box plots denote the distribution of cells with high ROS levels, and green box plots correspond to cells with low ROS levels. Each box represents the interquartile range (25th-75th percentiles), with the central line indicating the median, whiskers showing the data range, and individual points representing outliers. This visualization highlights material- and time-dependent variations in cellular oxidative response.

**Figure 6:**
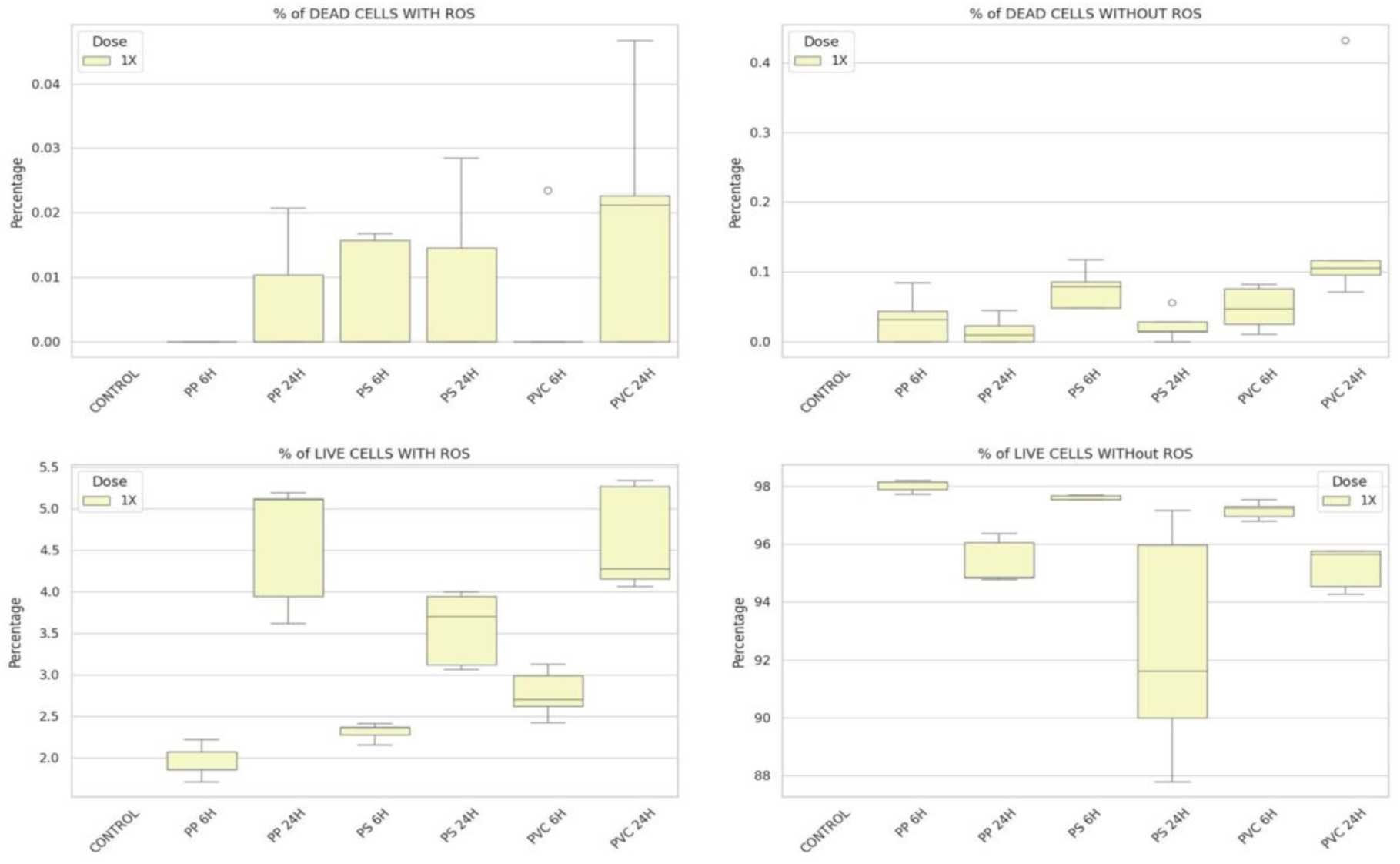
Distribution of cellular viability and oxidative stress status following NPs exposure. Box plots depict the percentage distribution of four cell populations: dead cells with ROS, dead cells without ROS, live cells with ROS, and live cells without ROS across control and NPs-exposed conditions. PBMCs were exposed to PS, PP, and PVC at a single dose (1X) and analyzed at 6 h and 24 h post-exposure. Control samples represent unexposed cells. Each panel corresponds to a distinct cell category, with box plots showing median values. Interquartile ranges and data dispersion. The figure illustrates time- and polymer-dependent shifts in cell viability and ROS-associated phenotypes following NPs exposure.

**Figure 7:**
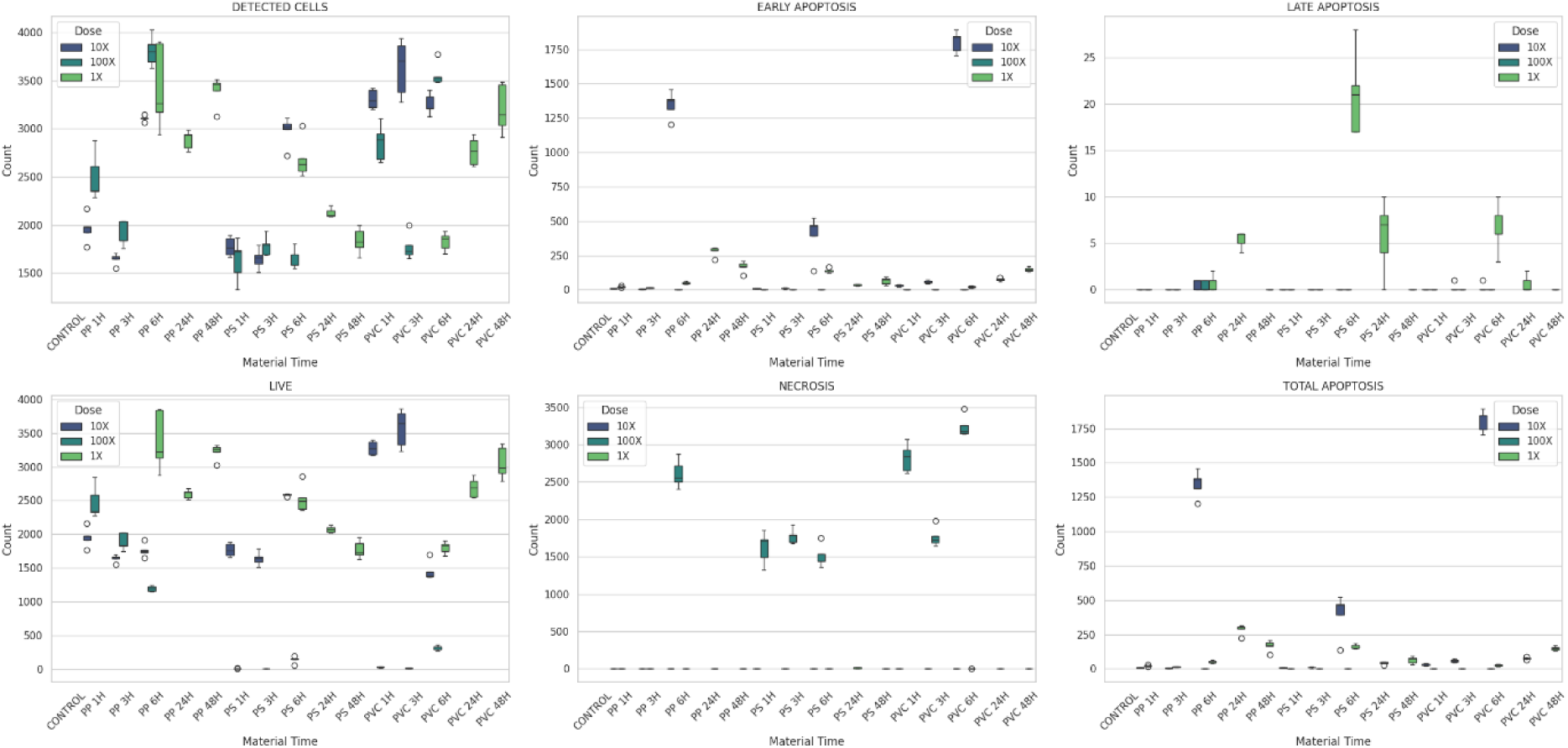
Distribution of cell fate outcomes following NPs exposure across materials, doses, and time points. Boxplots illustrate changes in total detected cells, early apoptosis, late apoptosis, live cell counts, necrosis, and total apoptosis after treatment with PS, PP, and PVC at 1X, 10X, and 100X concentrations for 6, 24, and 48 h.

**Figure 8:**
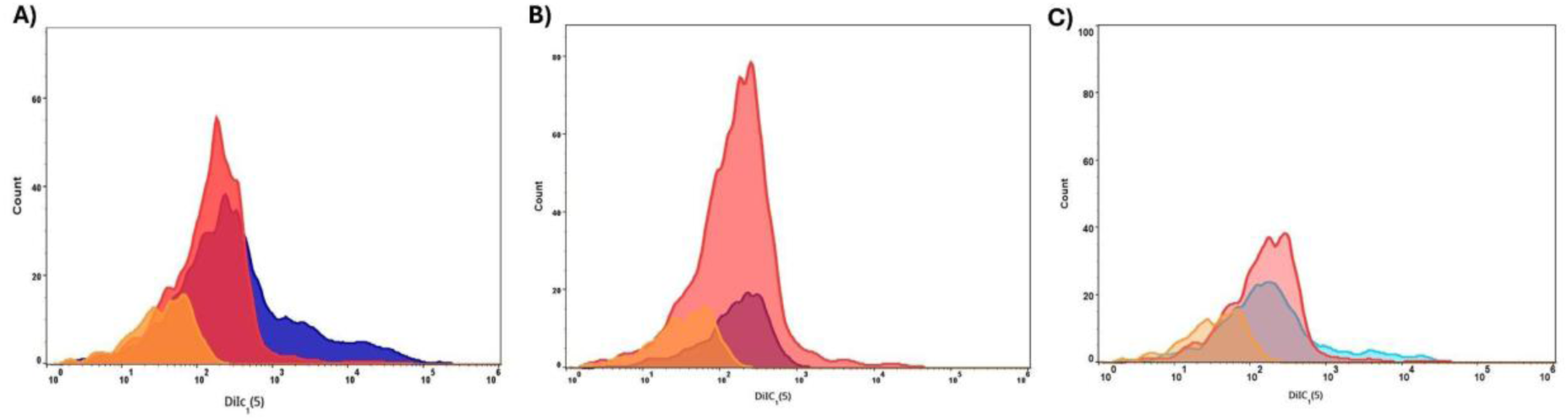
NPs exposure increases mitochondrial membrane potential as measured by DiIC1 (5) staining. **(A)** PP, **(B)** PS, and **(C)** PVC. Cells were exposed to each NPs type for 0 hr (orange), 6 hr (red), or 24 hr (blue). Plots display flow cytometry distributions of DiIC1 (5) fluorescence intensity (log scale, x-axis), where higher values represent higher mitochondrial membrane potential. For all NPs, the fluorescence peak shifts from left (control) to right (6 hr, 24 hr), indicating that mitochondrial membrane potential increases with duration of exposure.

### 3.3. Gene expression analysis

DRP1 was consistently upregulated in all NPs-treated groups relative to the control (**Figure 9A**). This elevation reflects impaired fission and indicates disrupted mitochondrial dynamics. Likewise, MFN1 showed marked upregulation upon NPs exposure **(Figure 9B**). PS-treated cells displayed the highest induction at 6 h, which declined gradually with prolonged exposure, while PP and PVC maintained moderately elevated expression levels. The results confirmed that NPs cause disturbances in the mitochondrial morphology and functioning, as alterations in the mitochondrial dynamics. To evaluate the impact of NPs on mitochondrial energy metabolism, expression patterns of ATP6, COX1, and ND6 were analyzed (**Figure 10**). ATP6 showed a time-dependent upregulation in PS-treated cells at 48 h, while PP and PVC induced minimal changes **(Figure 10A)**. COX1 was similarly upregulated in PS and to a lesser extent in PVC **(Figure 10B)**. ND6 expression was markedly elevated by PS, indicating strong activation of Complex I, whereas PP and PVC produced moderate increases **(Figure 10C)**, suggesting that NPs induce pronounced mitochondrial hyperactivation and elevated respiratory chain activity as an adaptive stress response. Analysis of BER genes revealed distinct transcriptional responses upon NPs exposure. OGG1 showed a rapid upregulation in PS-treated cells at 24 h, reflecting an immediate response to oxidative DNA damage, whereas PP and PVC induced moderate but prolonged activation **(Figure 11A)**. APE1 expression increased progressively over time, with the strongest induction under PVC treatment at 48 h, indicating sustained activation of DNA repair machinery in response to persistent oxidative stress **(Figure 11B)**. To assess mitochondrial stress signaling, the expression of OMA1 and DELE1 was examined **(Figures 12A and 12B)**. Both genes exhibited upregulation, suggesting that exposure to NPs induces ISR in the cell. Transcriptional profiling of DNA methyltransferases DNMT1, DNMT3a, and DNMT3b revealed significant suppression following NPs exposure, indicative of epigenetic disruption. DNMT1 expression declined progressively from moderate levels at 6 h to below at 48 h across all polymers **(Figure 13A)**. DNMT3a remained persistently low throughout all time points (**Figure 13B**). DNMT3b displayed transient material-specific levels, followed by a sharp decline at 48 h **(Figure 13C)**. Among the polymers, PP induced the most pronounced suppression, followed by PVC and PS (p < 0.01), reflecting polymer-specific epigenetic alterations.

**Figure 9:**
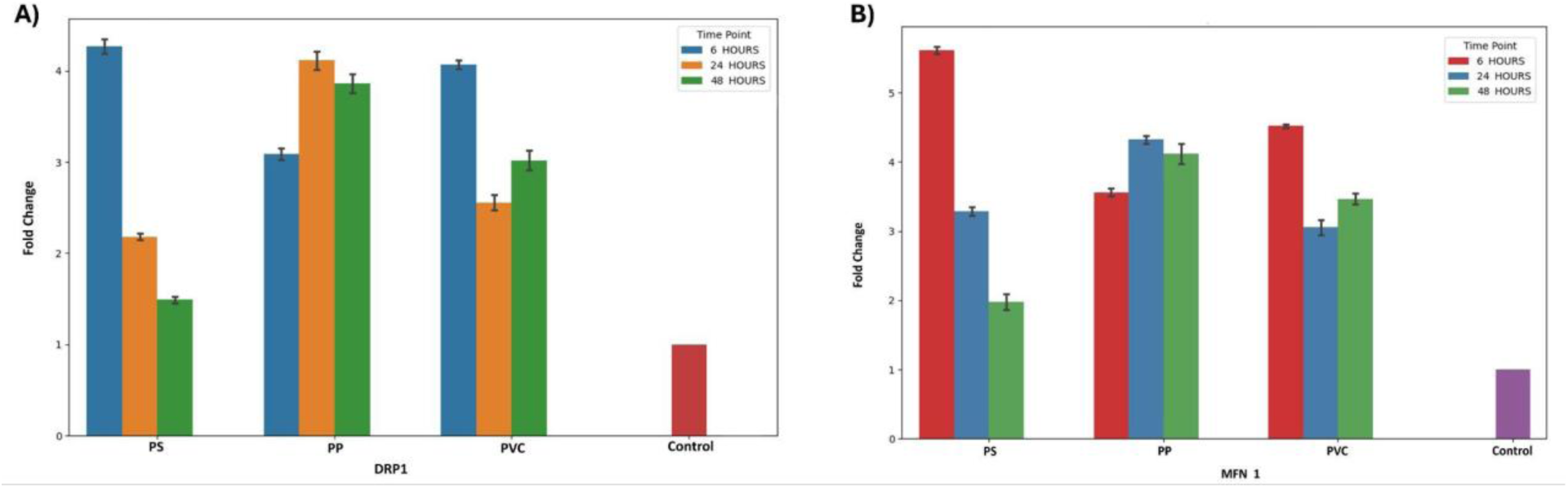
Fold change in **(A)** DRP1 and **(B)** MFN1 gene expression following NPs exposure at different time points. The figure presents quantitative changes in DRP1 and MFN1 expression relative to the untreated control, reflecting alterations in mitochondrial dynamics upon exposure to PS, PP, and PVC for 6, 24, and 48 h. Bars above a fold change of 1.0 denote gene upregulation, while values below 1.0 indicate downregulation, highlighting time- and material-dependent modulation of mitochondrial fission and fusion processes.

**Figure 10:**
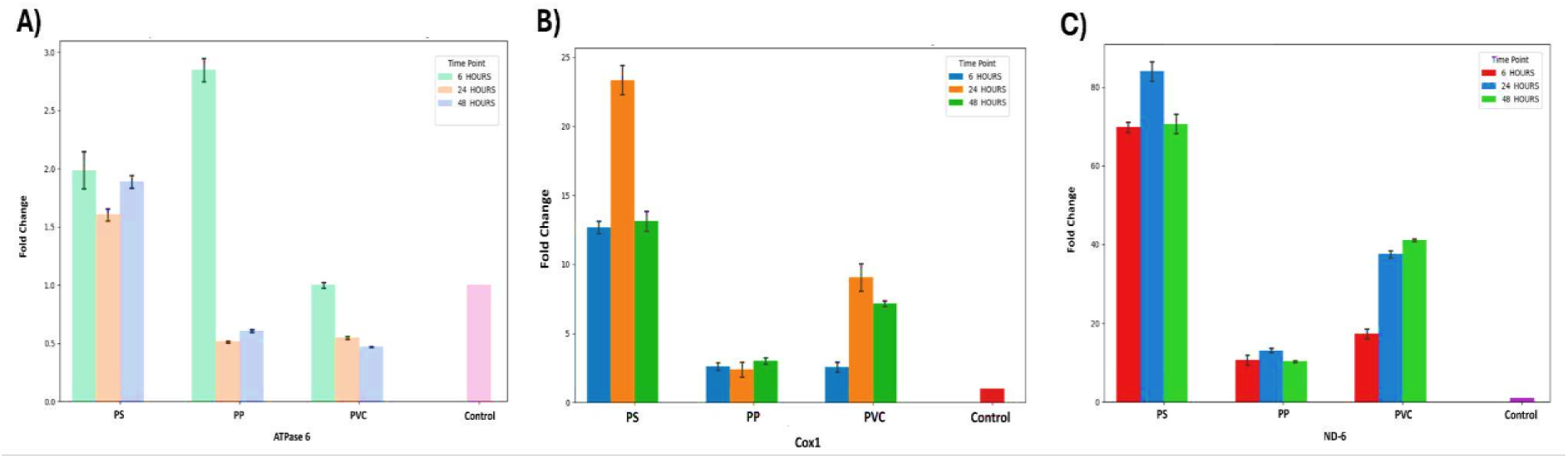
Expression levels in **(A)** ATPase 6, **(B)** COX1, and **(C)** ND6 following exposure to different NPs materials over time. The grouped bar chart depicts the relative expression levels of ATPase 6, COX1, and ND6 compared to untreated controls. The x-axis represents the mitochondrial genes, while the y-axis indicates the fold change in expression. Bars above 1.0 denote upregulation, and those below 1.0 indicate downregulation. Each color corresponds to a specific NPs type (PS, PP, or PVC) and exposure duration (6 h, 24 h, or 48 h). Notably, ATPase 6 exhibited pronounced upregulation under PS 24 h, PP 6 h, and PVC 48 h conditions, whereas COX1 and ND6 showed comparatively minor expression changes, indicating gene- and time-dependent responses to NPs exposure.

**Figure 11:**
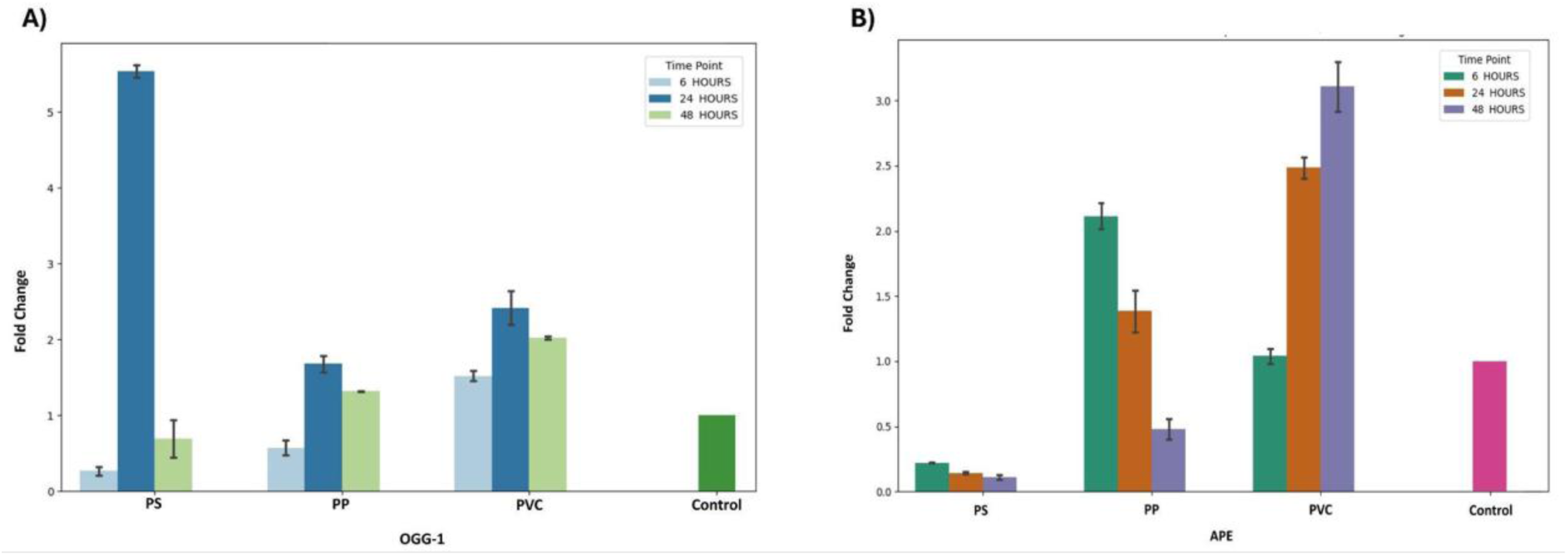
The figure illustrates relative expression levels of **(A)** OGG1 and **(B)** APE after exposure to different PS, PP, and PVC for 6, 24, and 48 h. Fold change values above 1.0 indicate upregulation, while those below 1.0 denote downregulation, highlighting time- and material-dependent modulation of mtDNA repair activity.

**Figure 12:**
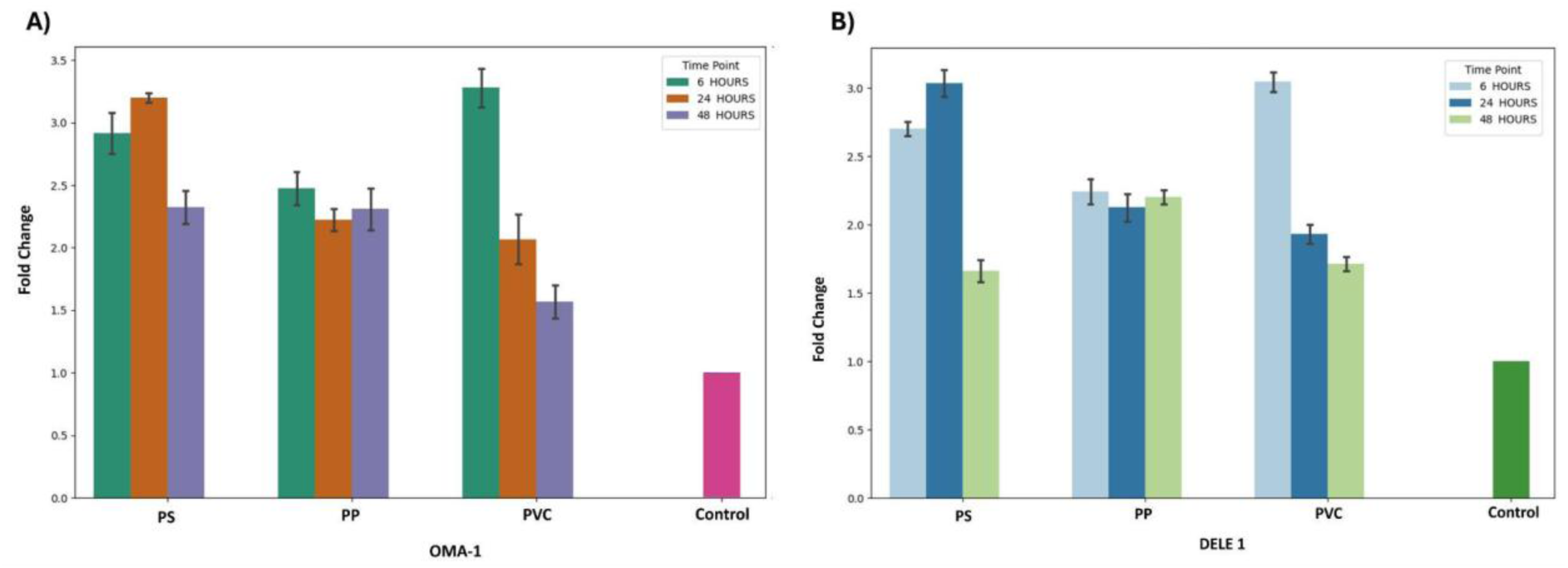
Mean fold-change expression of **(A)** OMA1 and **(B)** DELE1 in cells exposed to PS, PP, and PVC across 6, 24, and 48 h. OMA1 and DELE1 expression is strongly upregulated at 6 h, particularly in PS and PVC, but progressively declines at later time points. Control samples show minimal expression for both genes. Values represent mean ± SEM.

**Figure 13:**
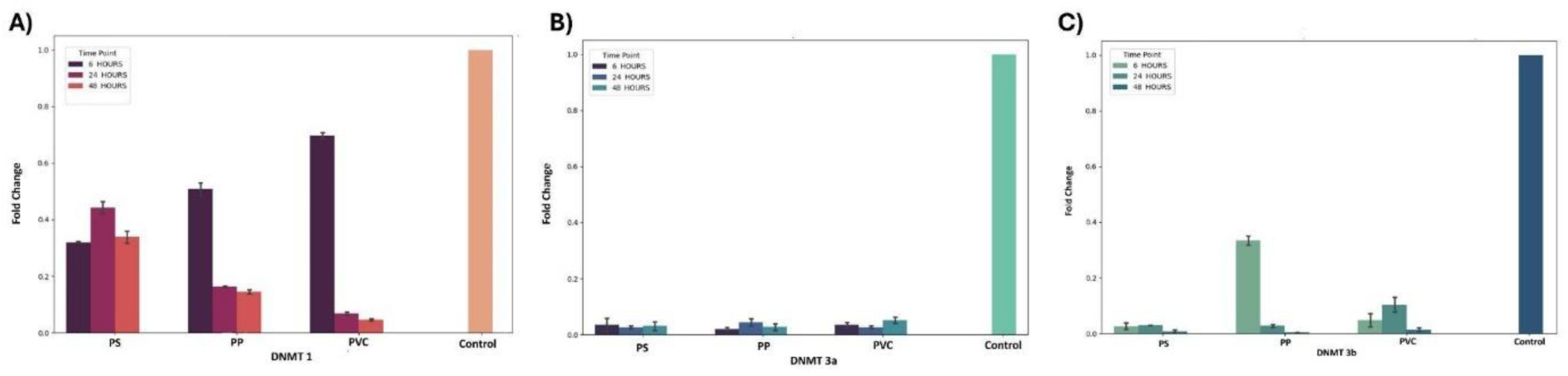
Fold change in DNA methylation gene expression following NPs exposure at different time points. This grouped bar chart depicts the fold change in expression of key DNA methylation genes, **(A)** DNMT1, **(B)** DNMT3a, and **(C)** DNMT3b, following exposure to different NPs types (PS, PP, and PVC) across 6, 24, and 48 h. The x-axis represents the individual genes, while the y-axis indicates relative fold change compared to the untreated control. Each color within a gene group corresponds to a specific NPs and exposure duration. Among all treatments, DNMT1 exhibited a striking upregulation, peaking under PVC exposure at 6 h (≈51-fold), suggesting a strong early epigenetic response, whereas DNMT3a and DNMT3b showed only minor expression changes across conditions.

Pearson correlation analysis revealed multiple statistically significant positive and negative associations among genes related to mitochondrial dynamics, oxidative phosphorylation, DNA repair, epigenetic regulation, and mitochondrial-associated microRNAs (p < 0.05). Strong positive correlations were observed between mitochondrial stress and dynamics markers, including DRP1 and OMA1 (r = 0.96, R² = 0.91, p < 0.001) and DRP1 and DELE1 (r = 0.98, R² = 0.95, p < 0.001). Epigenetic regulators also showed robust associations, with DNMT3a exhibiting a near-perfect positive correlation with DELE1 (r ≈ 1.00, R² ≈ 0.99, p < 0.001) and a strong correlation with DNMT3b (r = 0.85, R² = 0.73, p < 0.001). Among mitochondrial DNA–encoded oxidative phosphorylation genes, COX1 and ND6 were strongly positively correlated (r = 0.94, R² = 0.89, p < 0.001). Significant positive associations were additionally detected between ATPase 6 and FAPY (r = 0.70, R² = 0.49, p < 0.01), as well as between mitomiRs and mitochondrial markers, including miR-21 with FAPY (r = 0.69, R² = 0.48, p < 0.01) and miR-155 with ND6 (r = 0.36, R² = 0.13, p < 0.05). In contrast, significant negative correlations highlighted opposing regulatory relationships, notably between DRP1 and MFN1 (r = −0.54, R² = 0.29, p < 0.01) and between MFN1 and miR-34a (r = −0.64, R² = 0.41, p < 0.001). DNA repair genes also displayed inverse associations with mitochondrial genes, as APE was negatively correlated with COX1 and ND6 (r ≈ −0.42, R² ≈ 0.18, p < 0.05), while miR-21 showed a significant negative correlation with DNMT1 (r = −0.65, R² = 0.42, p < 0.01) (**Figure 14**).

**Figure 14:**
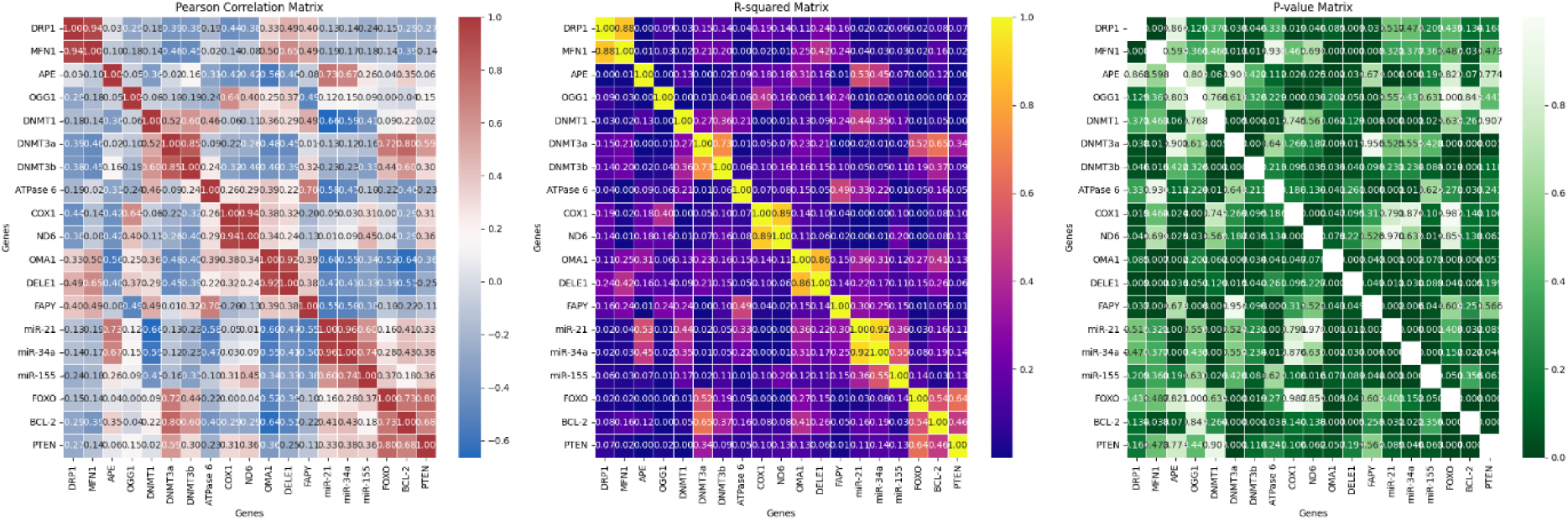
Gene Correlation Analysis. This figure provides a multi-panel visualization of the relationships among a set of genes and microRNAs (miRs). The Pearson Correlation Matrix (Left Panel) displays the strength and direction of the linear relationships (r), ranging from strong negative (blue) to strong positive (red). The R^2^ Matrix (Middle Panel) shows the coefficient of determination, indicating the proportion of variance explained between pairs (r^2^), with intense yellow representing the highest shared variance. Finally, the p-value matrix (right panel) presents the statistical significance of the correlations, where darker green indicates a lower p-value, suggesting a more statistically reliable correlation.

### 3.4 Alteration in DNA methylation

The methylation analysis revealed a clear and consistent hypomethylation across all examined mitochondrial regions (D-Loop1, D-Loop2, 12S, and 16S) in cells exposed to PS, PP, and PVC compared to the control. While the control group maintained baseline methylation, all NPs-treated conditions exhibited substantially reduced methylation levels, with the extent of hypomethylation varying depending on polymer type and exposure duration. PVC at 6 h showed the strongest hypomethylation, particularly in the D-loop regions, followed by PP and PS, indicating polymer-specific sensitivity. Across all areas, longer exposure did not restore methylation, indicating that NPs may persistently compromise mtDNA stability and function **(Figure 15)**.

**Figure 15:**
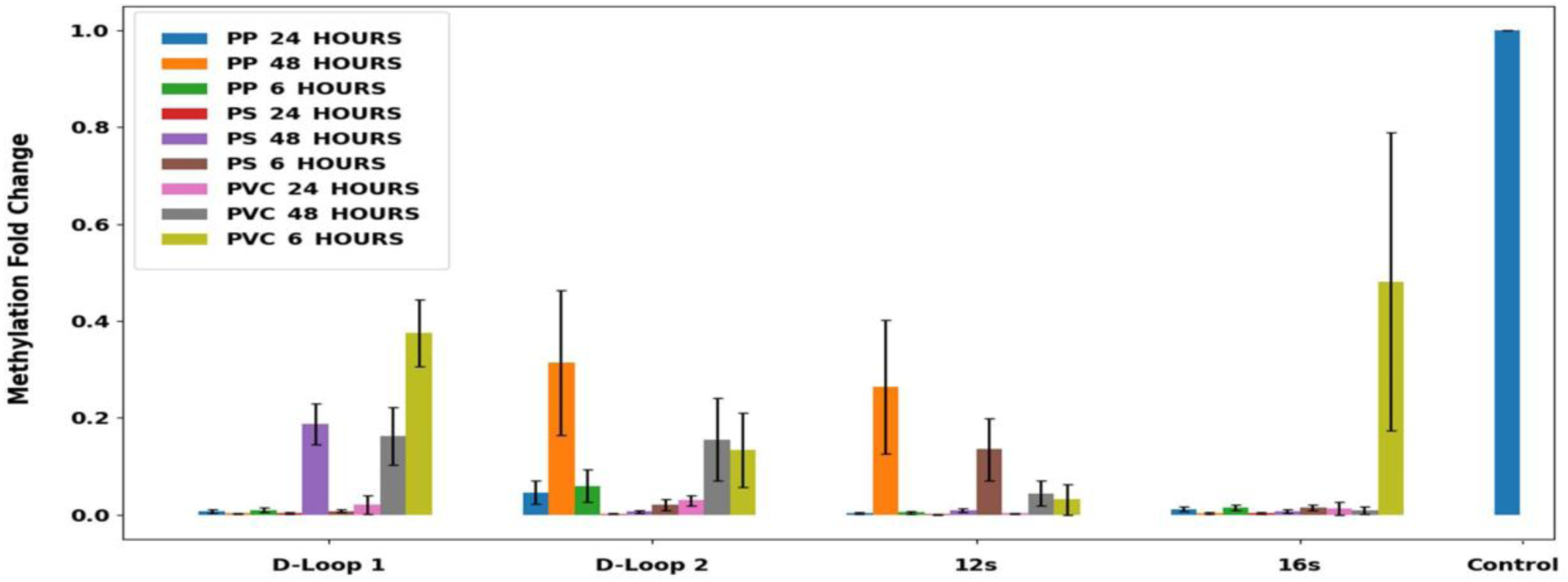
Methylation fold-change analysis of mitochondrial regions (D-Loop1, D-Loop2, 12S, and 16S) following exposure to PS, PP, and PVC NPs at 6, 24, and 48 h. All NPs-treated groups showed a marked reduction in methylation levels compared to the control (set to 1.0), indicating consistent hypomethylation across all regions. The degree of hypomethylation varied by polymer type and exposure duration, with PVC at 6 h showing the highest hypomethylation, followed by PP and PS. Error bars represent standard deviations (SD).

### 3.5 NPs exposure alters miRNA profiles and downstream target gene expression

PS, PP, and PVC induced a distinct time-dependent reprogramming of mitochondrial microRNAs (mitomiRs) **(Figure 16)**. All three mitomiRs, miR-21, miR-34a, and miR-155, showed pronounced early downregulation at 6 h, with PVC causing the strongest suppression. This inhibition persisted through 24 h, followed by a clear transition to upregulation at 48 h, where PVC again demonstrated the highest induction, followed by PS and PP. Correspondingly, the expression of downstream target genes Bcl2, FOXO3, and PTEN was altered in NPs-treated cells **(Figure 17)**, reflecting the functional impact of mitomiR modulation.

**Figure 16:**
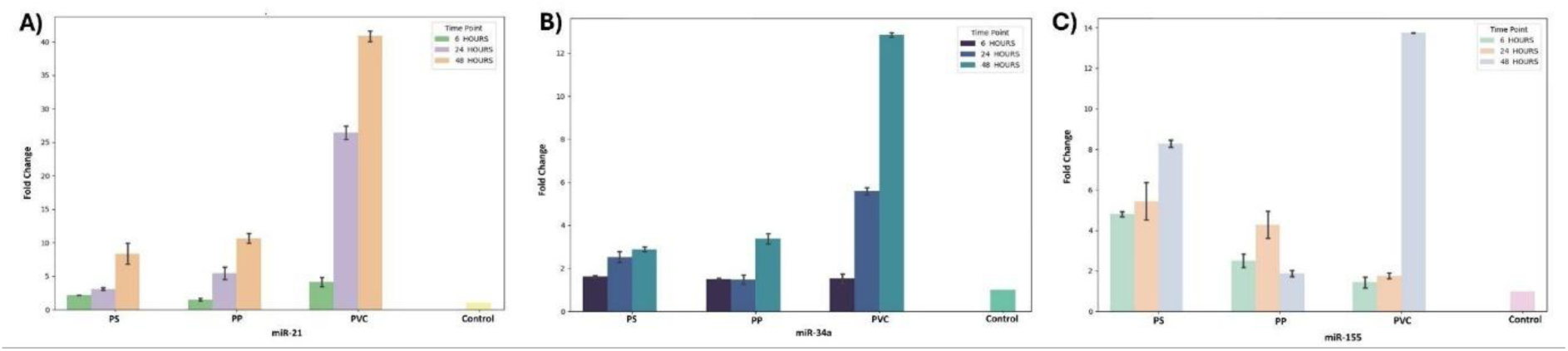
The bar graph represents the fold change in expression of three key miRNAs, **(A)** miR-21, **(B)** miR-34a, and **(C)** miR-155, following exposure to PS, PP, and PVC at 6, 24, and 48 h. The x-axis represents the miRNAs, while the y-axis shows their relative fold change compared to untreated controls. Each color denotes a specific NPs time combination. The results reveal substantial late-phase upregulation, with miR-21 showing a ∼106-fold increase under PVC exposure and miR-155 exhibiting a ∼108-fold rise under PS exposure at 48 h, indicating strong time- and material-dependent epigenetic activation in response to NPs exposure

**Figure 17:**
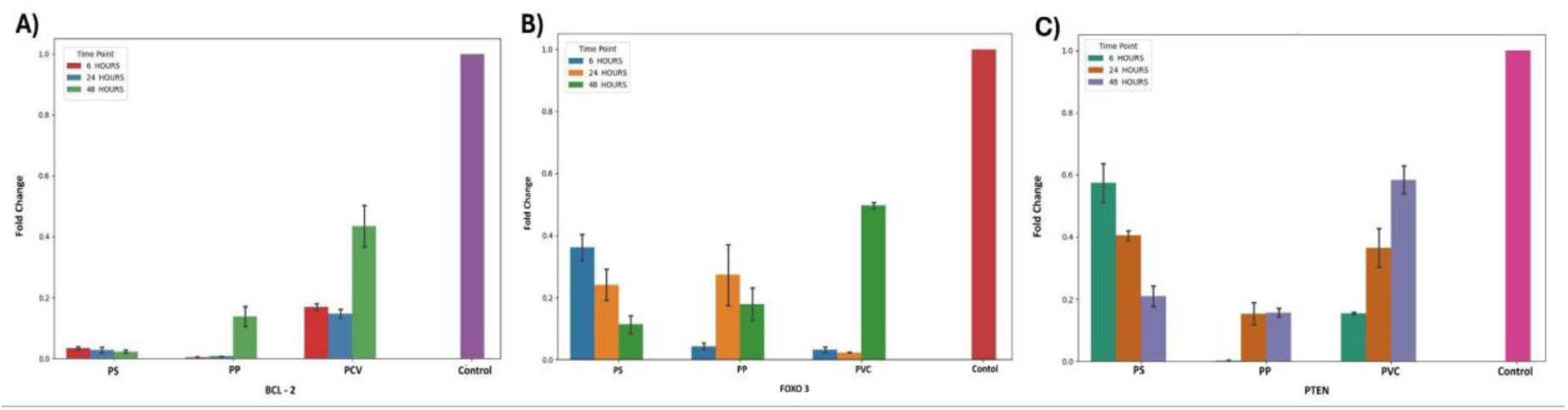
Mean expression levels of the miRNA target genes Bcl2, FOXO3, and PTEN in lymphocytes exposed to PS, PP, and PVC for 6, 24, and 48 h. Across all polymers and time points, NPs exposure consistently downregulated Bcl23, FOXO3, and PTEN compared with the control. **(A)** Bcl2 expression showed the strongest suppression under PVC at 48 h, indicating enhanced pro-apoptotic pressure. Similarly, **(B)** FOXO3 levels decreased progressively with time, particularly in PS- and PP-treated cells, reflecting compromised antioxidant and stress-response signaling. **(C)** PTEN expression also exhibited marked reduction across all NPs treatments, with PVC again eliciting the most pronounced decline at later time points, suggesting inhibition of key tumor-suppressive and regulatory pathways.

### 3.6 Evaluation of inflammatory cytokines

The expression profiles of key pro-inflammatory cytokines were evaluated in lymphocytes exposed to PS, PP, and PVC NPs for 6, 24, and 48 h. Exposure to PS, PP, and PVC induced distinct material and time-dependent alterations in the cytokine expression (**Figure 18**). IL-8 showed rapid and sustained upregulation, peaking at 48 h, with PVC eliciting the highest response (∼950 pg/mL), followed by PS and PP. This indicates a strong and persistent pro-inflammatory signaling loop throughout the exposure. TNF-α expression displayed a transient activation pattern, increasing at 6 h, peaking at 24 h, especially in PP-exposed cells, then declining by 48 h, returning near baseline in PVC-treated cells. This suggests a self-limiting inflammatory response. IL-6 levels rose at 6 h, with PP (∼580 pg/mL) and PS (∼500 pg/mL) showing the highest concentrations, while PVC elicited a moderate response (∼290 pg/mL). Elevated IL-6 persisted at 24 h, peaking in PP (∼610 pg/mL), before declining at 48 h, yet remaining above control levels. Correlation analysis indicated a strong positive association between TNF-α and IL-8 (r = 1.00), whereas IL-6 had minimal correlation with TNF-α (r = 0.16) and IL-8 (r = 0.14), indicating a more independent regulatory mechanism for IL-6.

**Figure 18:**
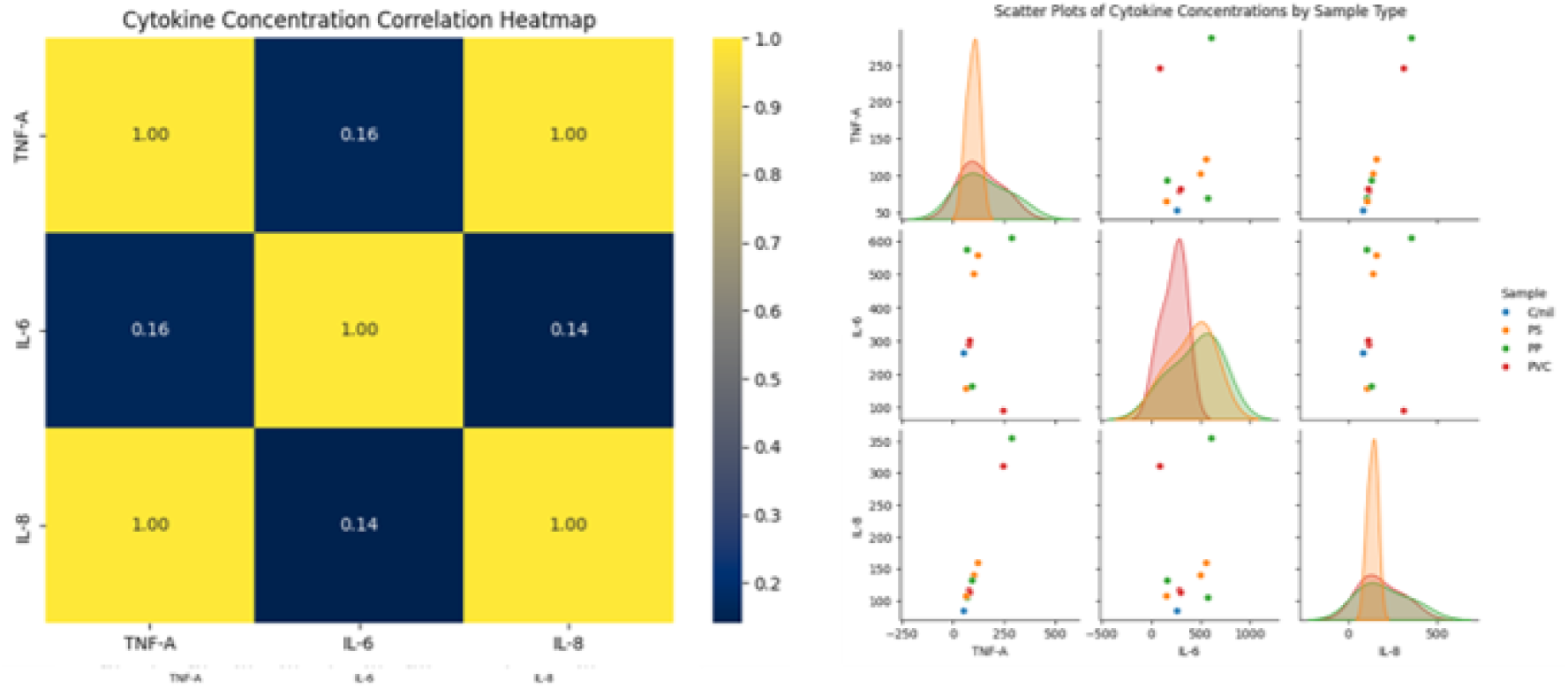
The figure presents a dual analysis of cytokine concentrations for TNF-α, IL-6, and IL-8. The left panel (heatmap) illustrates the Pearson correlation between these cytokines, notably showing a perfect correlation of 1.00 between TNF-α and IL-8, suggesting their concentrations track together across the samples, while the correlation of both with IL-6 is relatively weak (0.16 and 0.14, respectively). The right panel (Scatter Plots) further details these concentrations by sample type (Cnrl, PS, PP, PVC), using pairwise scatter plots to show relationships and diagonal Kernel Density Estimates (KDEs) to visualize the individual concentration distributions for each cytokine as they vary across the four distinct sample groups.

### 3.7 Measurement of mitochondrial electron chain complexes activity

NPs exposure induced distinct, polymer- and time-dependent alterations in mitochondrial respiratory complex activities (I-V). Complex I exhibited a consistent reduction across all treatments, with PVC and PP causing the strongest inhibition and PS showing comparatively preserved activity **(Figure 19A)**. Complex II demonstrated a transient early increase most pronounced in PVC and PP, followed by a decline, indicating short-lived compensatory enhancement of succinate dehydrogenase function **(Figure 19B)**. Complex III remained relatively stable, with minor fluctuations; PS and PP maintained slightly higher activity early on, whereas PVC induced a uniform decline **(Figure 19C)**. Complex IV displayed clear material-specific divergence: PS and PP induced early upregulation, while PVC caused a sustained decrease, suggesting impaired cytochrome c oxidase activity under persistent stress **(Figure 19D)**. Complex V activity increased initially across all treatments, particularly in PS and PVC, but declined at later time points, with PVC showing the greatest inhibition **(Figure 19E)**. Overall, these findings demonstrate that NPs disrupt oxidative phosphorylation in a progressive and polymer-dependent manner.

**Figure 19:**
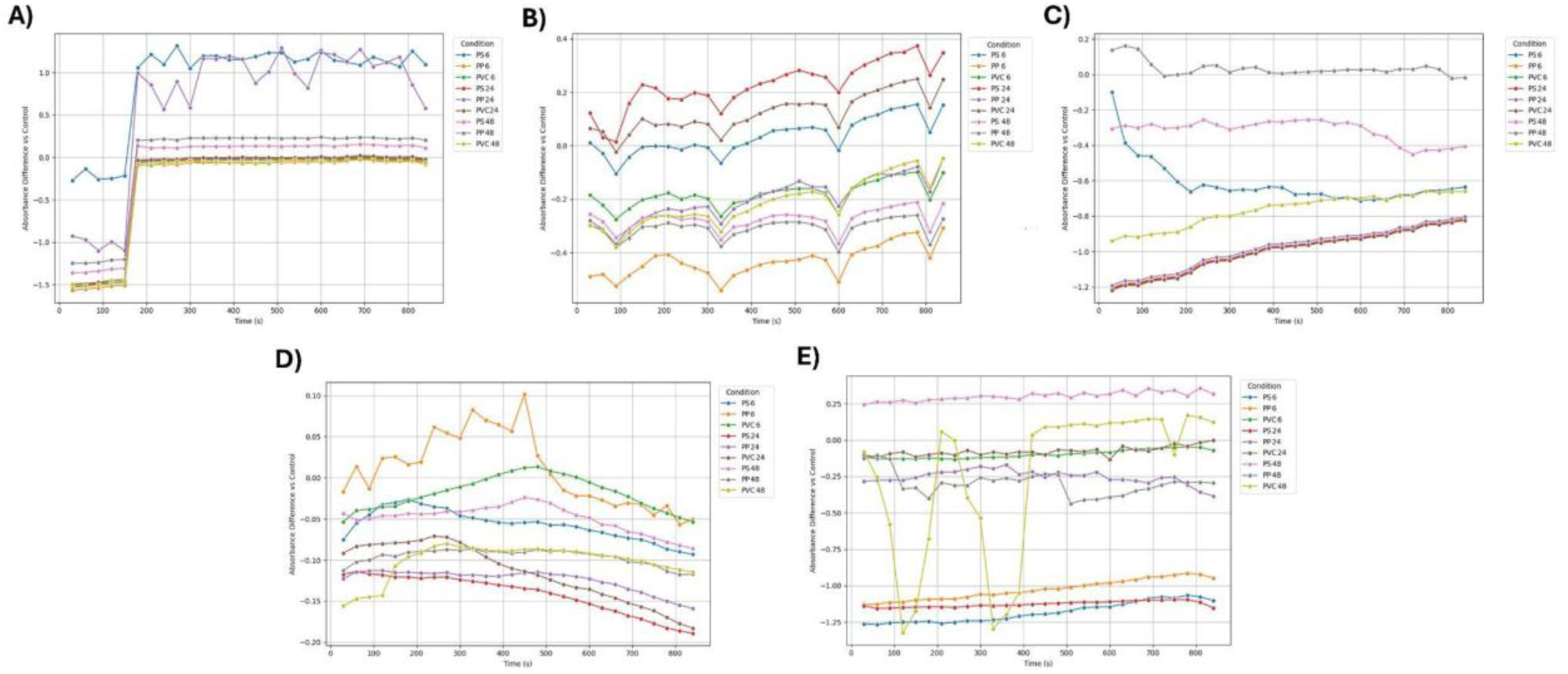
Each panel **(A-E)** represents one mitochondrial complex, comprising a series of five distinct kinetic line plots, labeled mitocomplex I - mitocomplex V, each illustrating the activity of a single mitochondrial respiratory chain complex over a time course. The y-axis represents the absorbance difference vs. control, quantifying the complex’s activity relative to a baseline, while the x-axis is time (s). Within each graph, multiple lines denote different experimental conditions, specifically the exposure of cells to various plastic types (PS, PP, PVC) for varying durations (6 h, 24 h, 48 h), thus allowing for a visual comparison of how each specific plastic treatment affects the kinetic activity of the individual mitochondrial complexes.

### 3.8 In silico molecular docking

Molecular docking analysis revealed distinct interaction profiles of oxidized NPs oligomers with mitochondrial complex I. Complex I exhibited the highest overall binding susceptibility, with docking scores ranging from - 9.558 to -6.074 kcal/mol. The strongest interaction (-9.558 kcal/mol) was observed for α-carboxyl-ω-hydroxyl poly(vinyl chloride) at the iron-sulfur (Fe-S) cluster of Complex I **(Figure 20A)**. The ligand’s carboxyl and hydroxyl groups formed hydrogen bonds with nearby polar residues. At the same time, its vinyl chloride backbone engaged non-polar residues through hydrophobic contacts, collectively stabilizing the binding and indicating a high potential for FeS-mediated electron transfer disruption. At the lipoyl-like cofactor site, polystyrene showed a similarly strong affinity (-9.238 kcal/mol) (**Figure 20B**). Notably, α-carboxyl-ω-hydroxyl poly(propylene) also demonstrated strong affinity for the Fe-S cluster, yielding a docking score of -8.897 kcal/mol, indicating substantial interaction potential comparable to PVC at this redox-active site **(Figure 20C)**. At the Fe-S cluster, carboxyl and hydroxyl functional groups facilitated hydrogen bonding with surrounding polar residues, while hydrophobic interactions between the polymer backbone and non-polar residues further stabilized ligand binding, suggesting a capacity for interference with Fe-S-mediated electron transfer. Overall, the docking results indicate that Complex I is substantially more vulnerable to NPs interaction, with the Fe-S cluster and lipoyl-like cofactor regions emerging as the most sensitive binding hotspots for oxidized NPs derivatives.

**Table 1.** Docking scores for oxidized NPs with complex I (5XTD)

| **Nanoplastics (Ligand)** | **Docking score (kcal/mol)** |
| --- | --- |
| **At Flavin mononucleotide (FMN) binding site** |  |
| a-Carboxyl-ω-hydroxypoly(styrene) | -7.99251 |
| a-Aldehyde-ω-hydroxypoly(styrene) | -6.75032 |
| PS Monomer terminal carboxyl | -6.66836 |
| a-Carboxyl-ω-hydroxypoly (vinyl chloride) | -6.15087 |
| PVC MONOMER 2 | -6.01624 |
| **At (NADPH) binding Site** |  |
| a-Carboxyl-ω-hydroxypoly(styrene) | -6.07418 |
| PS Monomer terminal carboxyl | -5.48992 |
| a-Carboxyl-ω-hydroxypoly(propylene) | -5.47423 |
| a-Aldehyde-ω-hydroxypoly(styrene) | -5.35733 |
| a-Carboxyl-ω-hydroxypoly(vinyl chloride) | -5.2027 |
| **At the iron-sulfur cluster binding site** |  |
| a-Carboxyl-ω-hydroxypoly(vinyl chloride) | -9.5581 |
| a-Carboxyl-ω-hydroxypoly(propylene) | -8.89698 |
| PVC MONOMER 2 | -7.21829 |
| a-hydroxyl-ω-hydroxypoly(vinyl chloride) | -4.23688 |
| poly(styrene) | -4.17244 |
| **At the lipoyl-like Cofactor binding site** |  |
| poly(styrene) | -9.23752 |

**Figure 20A:**
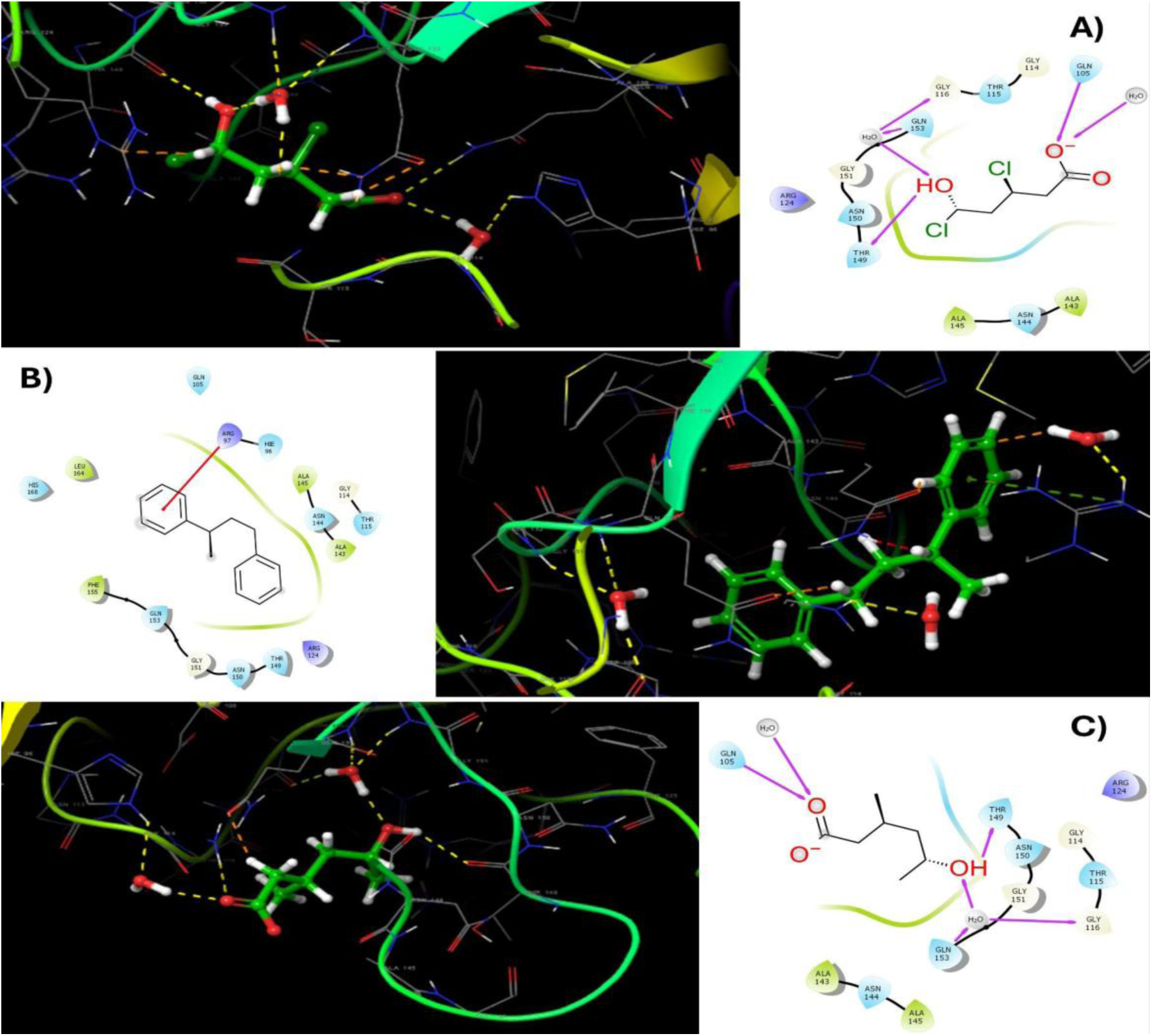
2D and 3D interaction diagrams of α-carboxyl-ω-hydroxyl poly(vinyl chloride) bound to the Fe-S cluster of mitochondrial complex I. This ligand showed the strongest affinity (-9.558 kcal/mol), fitting tightly within the Fe-S region and forming stabilizing hydrogen bonds and hydrophobic interactions indicative of potential interference with electron transfer. **Figure 20B**: 2D and 3D interaction diagrams of poly(styrene) docked at the lipoyl-like cofactor site of complex I. Polystyrene displayed strong binding (-9.238 kcal/mol), stabilized by hydrophobic and π–π interactions with surrounding aromatic residues. **Figure 20C**: 2D and 3D interaction diagrams of α-carboxyl-ω-hydroxyl poly(propylene) bound to the Fe-S cluster of mitochondrial complex I.

### 3.9 Robust predictive modeling of ND6 expression from mitocomplex I activity

The multi-output Random Forest regression model developed to predict ND6 gene expression from Mitocomplex I activity demonstrated strong and consistent performance across all evaluated targets when assessed on an independent test dataset. The model achieved uniformly high coefficients of determination (R²), ranging from approximately 0.984 to 0.987, indicating that more than 98% of the variance in ND6 gene expression was explained for all outputs, with PVC6 and PS24 showing the highest predictive accuracy **(Figure 21A)**. Correspondingly, mean squared error (MSE) values were low and narrowly distributed (approximately 0.014-0.017), reflecting minimal prediction error and stable performance across all ND6 endpoints **(Figure 21B)**. Residual analyses revealed symmetric residual distributions centered around zero with no systematic patterns, suggesting the absence of prediction bias or heteroscedasticity and confirming that the model effectively captured nonlinear relationships between Mitocomplex I activity and ND6 expression **(Figure 22)**. Furthermore, actual-versus-predicted plots showed a close alignment of predicted values with the identity line across the full dynamic range of expression levels, demonstrating strong agreement between PCR-quantified ND6 measurements and model predictions **(Figure 23)**. These outcomes display that the AI-based multi-output regression framework provides an accurate, robust, and generalizable approach for simultaneously predicting multiple ND6 gene expression targets from experimentally measured Mitocomplex I activity.

**Figure 21:**
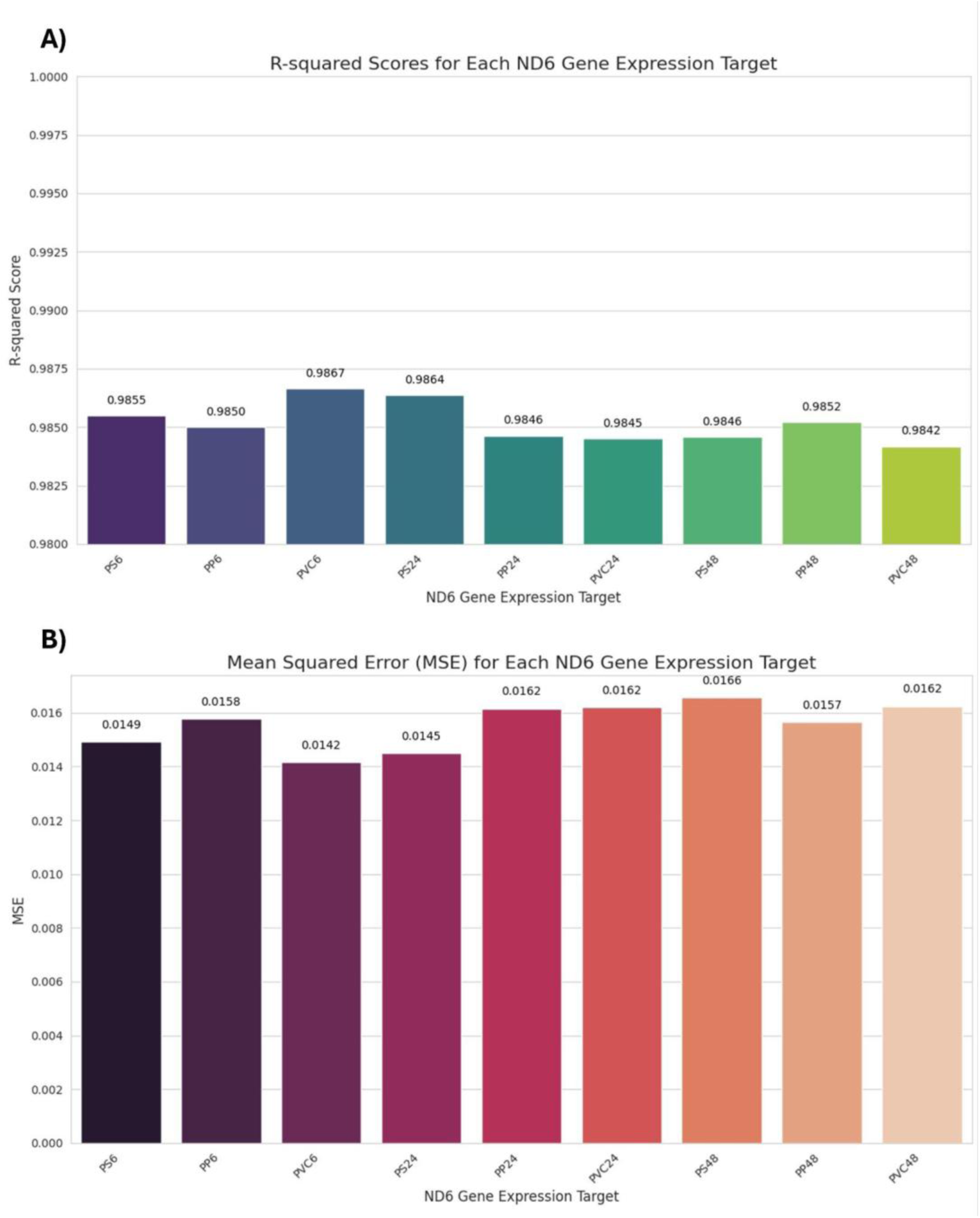
Model performance metrics for AI-based prediction of ND6 gene expression. **(A)** R-squared (R²) values for each ND6 gene expression target predicted from mitocomplex I activity using the multi-output Random Forest regression model. High and consistent R² values across all targets indicate strong explanatory power and uniform predictive performance. **(B)** Mean squared error (MSE) values for each ND6 gene expression target evaluated on the independent test dataset, demonstrating low and stable prediction errors across all outputs.

**Figure 22:**
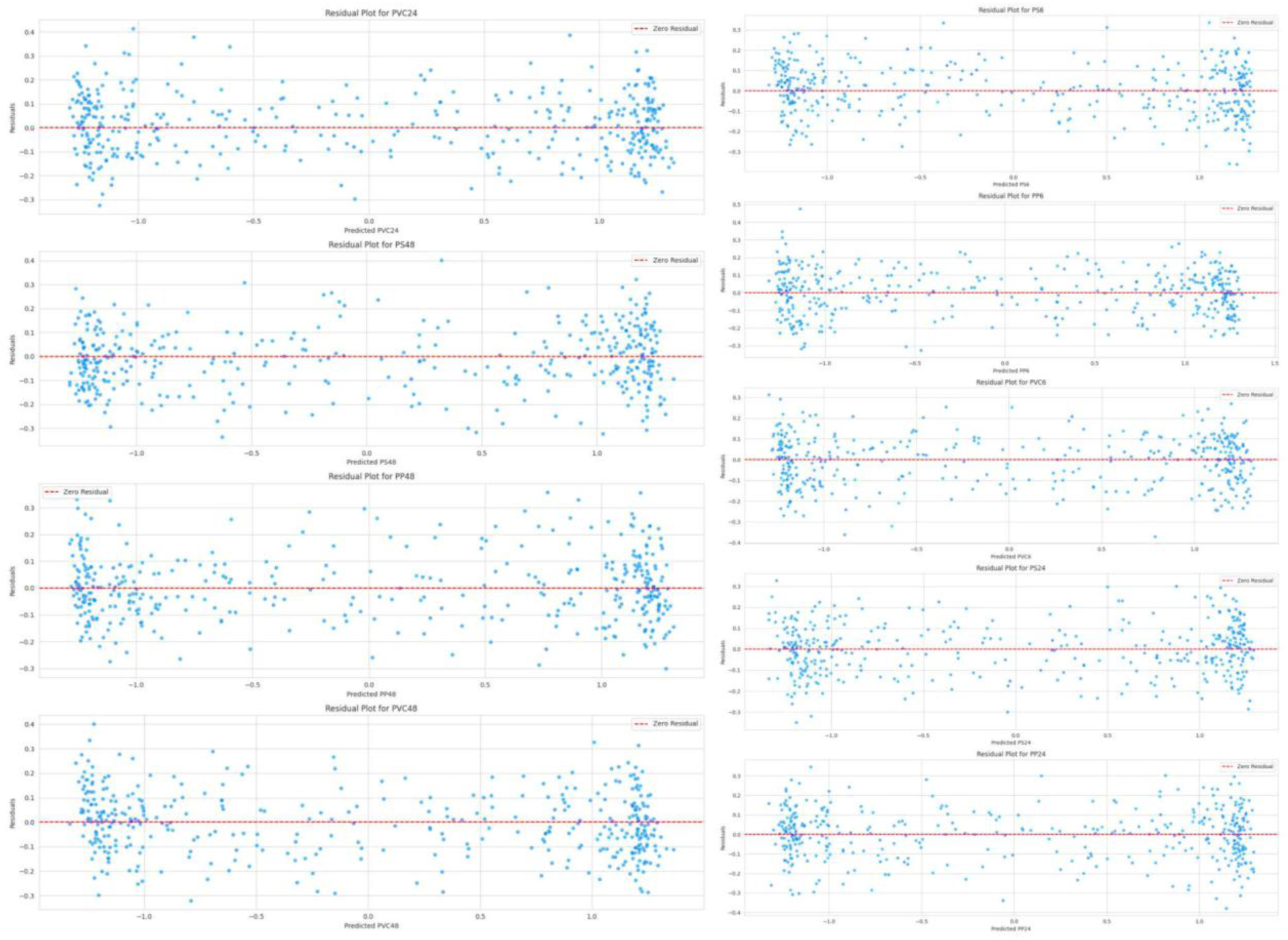
Residual analysis of ND6 gene expression predictions. Residual plots for each ND6 gene expression target showing the distribution of prediction errors as a function of predicted values. The dashed horizontal line represents zero residual. Residuals are symmetrically distributed around zero with no systematic trends, indicating minimal bias, a homoscedastic error structure, and adequate model fit across the full prediction range.

**Figure 23:**
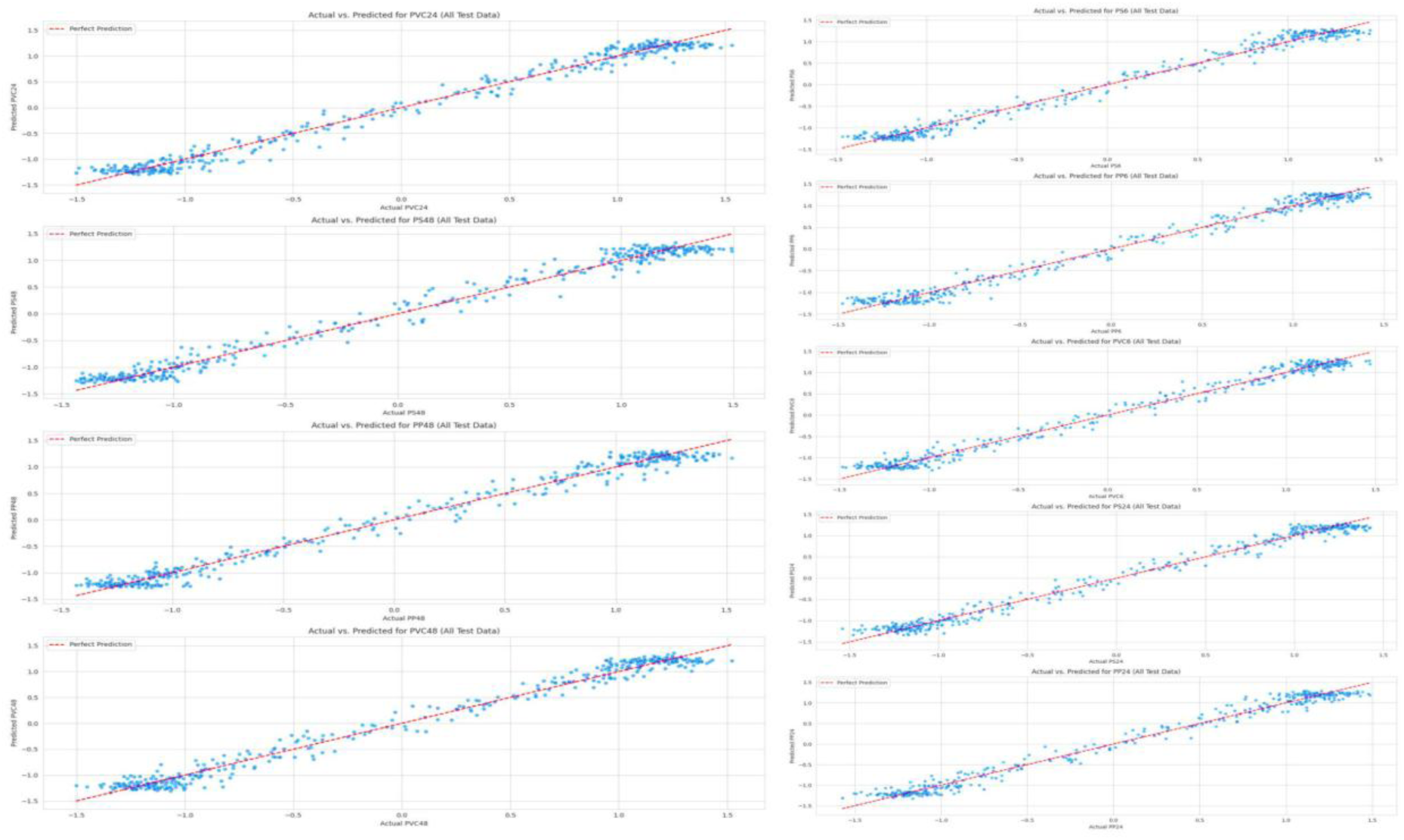
Actual versus predicted ND6 gene expression values. Scatter plots illustrate the relationship between experimentally measured ND6 gene expression levels, quantified by PCR, and the corresponding values predicted by the multi-output Random Forest regression model for all ND6 targets in the independent test dataset. The tight clustering of data points along this line across the full dynamic range of expression indicates a strong concordance between observed and predicted values, minimal systematic deviation, and effective model generalization.

## 4. Discussion

NPs are increasingly recognized as emerging immunomodulatory pollutants capable of disrupting multiple cellular pathways (Fan & Ha, 2025). The present study provides an integrated assessment of mitoepigenetic, redox imbalance, and immune-metabolic framing induced by three environmentally relevant NPs: PS, PP, and PVC across multiple exposure durations. By examining oxidative stress kinetics, mitochondrial dynamics, respiratory enzyme activities, gene expression changes, and computational docking interactions followed by AI prediction analysis. These findings collectively demonstrate that NPs exposure triggers polymer-specific oxidative stress, mitochondrial dysfunction, and downstream epigenetic and inflammatory alterations, highlighting mitochondria as central mediators of NPs toxicity. The physicochemical characteristics of polymer NPs significantly influence their colloidal stability and interactions with biological systems. DLS analysis showed that PS exhibited the smallest Z-average hydrodynamic diameter, and the others are also in the nano range (Gurjar et al., 2024). Low electrostatic repulsion of all three NPs suggests interactions are dominated by hydrophobic and van der Waals forces, favoring membrane and protein association. Polymer-specific size differences may influence cellular uptake and mitochondrial interactions (Bhattacharjee et al., 2012). PS were, therefore, selected for cellular uptake studies due to their smaller size. DIC fluorescence microscopy demonstrated time-dependent internalization of PS particles by lymphocytic cells, with minimal nonspecific association at baseline and punctate intracellular fluorescence observed after 1 h, indicating active uptake. Increased fluorescence intensity and widespread cytoplasmic distribution at later time points reflected sustained internalization. Flow cytometric analysis corroborated these findings, revealing a progressive rightward shift in fluorescence intensity with increasing exposure duration. Thus, the efficient internalization of NPs by immune cells raises concerns regarding potential disruption of cellular homeostasis and immune signaling, underscoring possible immunotoxicological risks associated with environmental support. NPs are clinically immunomodulatory particles capable of entering systemic circulation and directly engaging immune cells and further disrupting mitochondrial integrity (Liu et al., 2022). NPs induced a surge in Fpg-sensitive oxidized purines at 6 h, most pronounced with PP, indicating acute ROS-driven purine damage preceding effective repair or antioxidant compensation (Li et al., 2023). The decline in FAPY activity at 24-48 h, despite sustained cytosolic and mtROS, indicates exhaustion of BER capacity and preferential loss of severely damaged cells. This redox repair imbalance is consistent with flow cytometric evidence showing persistent mtROS accumulation, mitochondrial membrane depolarization, and altered cell fate decisions. PP sustains prolonged mtROS, ΔΨm collapse, and necrotic cell death, whereas PS and PVC induce transient oxidative bursts followed by delayed activation of apoptotic pathways (Yöntem & Ahbab, 2024).

Mitochondrial structural and genetic indicators further confirmed NPs exposure vulnerability. PS increased DRP1 indicates enhanced mitochondrial fission in response to stress, while elevated MFN1 suggests a compensatory rise in mitochondrial fusion to preserve mitochondrial integrity. PS exposure elicited the most pronounced early changes, particularly at 6 h, indicating rapid mitochondrial sensing and adaptation, whereas PP and PVC produced sustained but comparatively moderate effects. These changes reflect the cell’s attempt to balance fission-fusion dynamics and restore mitochondrial homeostasis under NPs-induced oxidative stress (Chang et al., 2023). Likewise, the biphasic modulation of mitochondrial-encoded genes (MT-ATP6, MT-COX1, and MT-ND6) indicated an initial adaptive enhancement of OXPHOS gene expression; PS strongly upregulated ATP6, COX1, and particularly ND6, while PP and PVC induced transient transcriptional activation of these genes, accumulating oxidative load, consistent with mitochondrial stress responses (Bennet et al., 2021). Because mitochondrial metabolites regulate epigenetic enzymes, potentially contributing to chronic inflammation and metabolic dysregulation (Bhargava et al., 2018). The elevated expression of DNA repair markers in all NPs more pronounced in PS in OGG1, while APE expression by PVC points to the activation of the BER pathway (Singh et al., 2017), reflecting oxidative base damage in mitochondrial genomes (Mishra et al., 2022b). In parallel, upregulated OMA1 and DELE1 expression suggests that NPs exposure may induce a pro-survival signaling mechanism inside the host cell through the activation of the ISR. Altogether, these outcomes showcase that NPs exposure can trigger an integrated cellular stress network, implying that even low doses at prolonged durations may facilitate cumulative mitochondrial strain progressively. The marked suppression of DNMT1, DNMT3a, and DNMT3b expression following NPs exposure highlights a significant disruption of the cellular epigenetic machinery. Reduced DNMT activity is consistent with the observed hypomethylation across key mitochondrial genomic regions, including D-loop1, D-loop2, 16S rRNA, and 12S rRNA, which are critical for mitochondrial integrity (Mishra et al., 2022a; Liu et al., 2025); such loss of methylation can impair mtDNA stability. The differential changes in mitochondrial respiratory complex activities further underscore the material-specific effects of NPs on OXPHOS, with sustained Complex I inhibition, most pronounced in PVC- and PP-exposed cells, indicating early electron transport disruption and increased vulnerability to oxidative damage, whereas PS showed relative enzymatic preservation despite transcriptional hyperactivation. NPs selectively disrupt Complex I, leading to mitochondrial injury and reduced ATP production. The transient increases in Complex II and initial elevations in Complexes IV and V likely represent short-lived compensatory mechanisms aimed at maintaining ATP synthesis under oxidative stress, which subsequently dwindle complex activities over time. The relatively stable but fluctuating response of Complex III further underscores the selective sensitivity of individual mitochondrial complexes to NPs-induced dysfunction (Huang et al., 2023). PVC’s consistent induction of the most severe and sustained reductions in complex activities reflects its potent disruptive influence on mitochondrial bioenergetics. The mitoepigenetic dimension was further supported by modulation of mitomiRs (miR-21, miR-34a, miR-155). In the present study, NPs-treated cells showed elevated expression of mitomiR-21, mitomiR-34a, and mitomiR-155. This alternation in miRNA was further confirmed by observing the downregulation pattern in its respective target genes, Bcl2, FOXO3, and PTEN. These coordinated mitomiRs target gene interactions highlight a correlative mitoepigenetic axis through which NPs exposure drives mitochondrial dysfunction and cytotoxicity via integrated transcriptional and post-transcriptional regulation (Chattopadhyay et al., 2024). Correlation analyses reveal mitochondrial Complex I as the primary vulnerability in NPs-exposed cells. Positive correlations between DRP1 and the stress effectors OMA1 and DELE1 indicate coordinated activation of mitochondrial fission and retrograde signaling, consistent with impaired electron transfer at Complex I (Lee et al., 2022; Huang et al., 2023). The strong association between DNMT3a and DELE1 further couples mitochondrial stress to epigenetic remodeling. Coordinated upregulation of mtDNA-encoded OXPHOS genes, including MT-ND6, suggests compensatory transcriptional responses to Complex I dysfunction (Lin et al., 2022). Conversely, negative correlations between DRP1 and MFN1 reflect a shift toward fission-dominated dynamics, suppressing fusion. The inverse relationship between MFN1 and miR-34a supports microRNA-mediated repression of fusion pathways, promoting mitochondrial fragmentation (Chattopadhyay et al., 2024). The negative correlations between the DNA repair enzyme APE1 and Complex I-associated genes highlight a trade-off between bioenergetic demands and mtDNA maintenance under elevated oxidative stress from Complex I impairment (Xu et al., 2021). Moreover, our findings align with mechanistic evidence through which NPs can activate inflammatory responses through ROS-mediated MAPK and NF-κB signaling (Liu et al., 2020). The confirmed elevation of IL-6, TNF-α, and the sustained upregulation of IL-8 support this ROS-dependent inflammatory cascade (Bhargava et al., 2019), particularly in PVC-exposed cells (Chen et al., 2023). With respect to cytokine kinetics, IL-6 peaks under early PP and PS exposure, while IL-8 induction remains persistent, suggesting differential temporal modulation of inflammatory outputs, potentially reflecting distinct ROS dynamics and downstream transcriptional regulation. The observed strong correlation between TNF-α and IL-8 reinforces shared NF-κB regulation, whereas the dissociated pattern of IL-6 implies additional layers of control, including mitochondrial or epigenetic influences. This study demonstrates that NPs exposure perturbs core cellular homeostasis by targeting mitochondrial machinery, leading to oxidative stress amplification, epigenetic reprogramming, and inflammatory activation. These convergent effects compromise bioenergetic stability and immune regulation, exerting sustained cellular impairments.

Computational docking analyses corroborated experimental observations by identifying mitochondrial Complex I as the primary and most functionally relevant interaction hub for oxidized NPs oligomers. High-affinity binding of oxidized PS, PP, and PVC oligomers was consistently localized near Fe-S clusters and lipoyl-like regions of Complex I, structural domains critical for electron transfer and proton translocation. Such interactions are likely to disrupt electron flow within the respiratory chain, thereby increasing electron leakage and predisposing mitochondria to excessive ROS generation. Integration of docking-derived structural insights with AI predictions revealed a strong functional coupling between Complex I impairment and ND6 gene expression. ND6, a mitochondrially encoded subunit integral to Complex I assembly and electron transport stability, is highly sensitive to redox imbalance and structural perturbation of the complex. The AI model’s ability to accurately predict ND6 expression from mitocomplex I activity indicates that NPs-induced disruption at the Complex I level is directly translated into transcriptional dysregulation of ND6. Oxidative functionalization of NPs oligomers (e.g., -COOH and -CHO groups) further amplified these effects by enhancing hydrogen bonding and protein-binding compatibility with Complex I domains, consistent with evidence that environmental aging increases NPs bioactivity and toxicity. Polymer-specific differences were also evident: PP induced a more sustained oxidative burden, while PVC elicited stronger inflammatory signaling, likely reflecting differences in binding stability and interaction geometry within Complex I.

The AI-based multi-output Random Forest regression model demonstrated strong and consistent performance (Soni et al., 2025a) in predicting ND6 gene expression from mitocomplex I activity, indicating a robust relationship between mitochondrial functional activity and ND6 transcriptional output. High and uniform R² values across all ND6 targets, together with low MSE scores, confirm that the model accurately captured the variance in gene expression while maintaining stable predictive performance across multiple endpoints. Residual and actual-versus-predicted analyses further indicated minimal bias, absence of systematic error, and effective generalization to unseen data. Methodologically, the use of a random forest regressor within a multi-output framework proved well-suited to this biological dataset, enabling simultaneous prediction of multiple ND6 expression targets while accommodating nonlinear relationships and the experimental variability inherent in ELISA and PCR-based measurements. Biologically, the strong predictive linkage supports the close coupling between Mitocomplex I activity and ND6 gene expression, consistent with the role of ND6 as a mitochondrially encoded component of Complex I. While external validation and deeper interpretability analyses would further strengthen these findings, the results demonstrate the utility of AI-driven modeling as a reliable tool for integrating mitochondrial functional assays with gene expression profiling. These outcomes bespeak Complex I as the central mechanistic nexus linking NPs-protein interactions, AI-predicted ND6 dysregulation, and downstream ROS-driven mitochondrial toxicity.

## Conclusion

NPs act as potent immunometabolic disruptors, primarily targeting mitochondrial function. Exposure to PS, PP, and PVC induced oxidative stress, mitochondrial dysfunction, selective inhibition of respiratory complex I, epigenetic alterations, and persistent inflammatory activation, establishing mitochondria as the primary mediator of NPs toxicity. Integrative analysis combining experimental data, computational docking, and AI-driven modeling revealed a key complex I-ND6 regulatory axis that mechanistically connects NPs-protein interactions to redox imbalance and mitochondrial gene dysregulation. PVC elicited the most pronounced mitochondrial membrane disruption and epigenetic changes; PP drove sustained oxidative stress and necrotic features; and PS promoted compensatory upregulation of mitochondrial gene expression, highlighting polymer-dependent modulation of mitochondrial homeostasis. However, limitations include reliance on an ex vivo PBMCs model, which may not capture tissue-specific or systemic dynamics; absence of long-term in vivo validation; and incomplete mimicry of real-world chronic, low-dose exposures to heterogeneous, environmentally aged NPs mixtures. Future studies should prioritize physiologically relevant exposure models, longitudinal organ-specific in vivo investigations, incorporation of aged and mixed NPs, and multi-omics integration with explainable AI to refine dose-response relationships and enhance human health risk assessments for these emerging NPs contaminants.

## Ethical Declaration

The study received approval from the Institutional Ethics Committee (IEC) of ICMR-NIREH, Bhopal (NIREH/BPL/IEC/2025-26-208).

## Acknowledgements

The authors are highly thankful to Mr. Sagar Patel for his technical assistance.

## Author Contributions (CRediT taxonomy)

PKM: Conceptualization, Methodology, Supervision, Project administration, Funding acquisition.

AKR, AC, AP, and RPT: Investigation and experimental execution.

VG and AA: Sample collection and physicochemical characterization.

VG and AKR: Data visualization and figure design.

VG, DKS, and AC: In silico molecular docking analyses.

RT and RKS: Data analysis and editing

AKR, AC, VG, AA, and PKM: Original draft preparation.

## Data availability

Data will be made available on request.

